# A versatile *in situ* cofactor enhancing system for meeting cellular demands for engineered metabolic pathways

**DOI:** 10.1101/2023.01.08.523081

**Authors:** Juthamas Jaroensuk, Chalermroj Sutthaphirom, Jittima Phonbuppha, Wachirawit Chinantuya, Chatchai Kesornpun, Nattanon Akeratchatapan, Narongyot Kittipanukul, Kamonwan Phatinuwat, Sopapan Atichartpongkul, Mayuree Fuangthong, Thunyarat Pongtharangkul, Frank Hollmann, Pimchai Chaiyen

**Affiliations:** School of Biomolecular Science and Engineering, Vidyasirimedhi Institute of Science and Technology (VISTEC); Wangchan Valley, Rayong, Thailand; Department of Biochemistry and Center for Excellence in Protein and Enzyme Technology, Faculty of Science, Mahidol University; Bangkok, Thailand; Program in Applied Biological Sciences, Chulabhorn Graduate Institute, Bangkok, Thailand; Laboratory of Biotechnology, Chulabhorn Research Institute; Bangkok, Thailand; Department of Biotechnology, Faculty of Science, Mahidol University; Bangkok, Thailand; Department of Biotechnology, Delft University of Technology; Van der Maasweg 9, Delft, 2629 HZ, Netherlands

## Abstract

Cofactor imbalance obstructs the productivities of metabolically engineered cells. Herein, we employed a minimally perturbing system, xylose reductase and lactose (XR/lactose), to increase levels of a pool of sugar-phosphates which are connected to the biosynthesis of NAD(P)H, FAD, FMN and ATP in *Escherichia coli*. The XR/lactose system could increase the amounts of the precursors of these cofactors and was tested with three different metabolically engineered cell systems (fatty alcohol biosynthesis, bioluminescence light generation and alkane biosynthesis) with different cofactor demands. Productivities of these cells were increased 2-4-fold by the XR/lactose system. Untargeted metabolomic analysis revealed different metabolite patterns among these cells; demonstrating that only metabolites involved in relevant cofactor biosynthesis were altered. The results were also confirmed by transcriptomic analysis. Another sugar reducing system (glucose dehydrogenase, GDH) could also be used to increase fatty alcohol production but resulted in less yield enhancement than XR. This work demonstrates that the approach of increasing cellular sugar phosphates can be a generic tool to increase *in vivo* cofactor generation upon cellular demand for synthetic biology.

**Teaser:** Use of sugar and sugar reductase to increase sugar phosphates for enhancing *in situ* synthesis of cofactors upon cellular demand for synthetic biology.

## Introduction

Synthetic biology and metabolic engineering provide greener solutions for the production of valuable chemicals compared to chemical-based approaches due to their less detrimental effects on the environment (*1, 2*). The approach of using microbial cell factories is also sustainable because it can convert renewable biomass (rather than petrochemicals or other non-renewable resources) into products of interest by fermentations (*1, 3, 4*). Creation of microbial cell factories requires strategic selection of enzymatic reactions and overexpression of their genes *in vivo*. To maintain good productivity, the relevant enzymes require a sufficient supply of substrates or cofactors^1^ e.g., NAD(P)H, FAD, FMN, ATP, acetyl-CoA) (*5-8*). Subsequently, the productivity of metabolically engineered cells is frequently hampered by the scarcity of substrates or cofactors required for enzymatic reactions owing to the limited amounts of these compounds available *in vivo*.

A well-established approach to overcome the insufficient supply of cofactors in microbes is incorporation of extra metabolic pathways for cofactor regeneration (*9, 10*). For example, formate dehydrogenase (FDH) or glucose dehydrogenase (GDH) are incorporated into cells to enhance generation of NAD(P)H from NAD(P)^+^ using formate and D-glucose as reductants (*11, 12*), while polyphosphate kinase (PPK) is used to regenerate ATP from ADP and polyphosphate (*5, 13*). Simple addition of substrates and cofactors (particularly charged molecules) is typically sub-optimal as they generally have low cell permeability and require further optimization such as adding a surfactant (*5*). Although this approach can somewhat enhance regeneration of selected cofactors, it is not entirely efficient because the total amount of cofactors i.e., NAD(P)H plus NAD(P)^+^ or ADP plus ATP does not change. Besides, biocatalytic systems often require more than one type of cofactors. Developing effective and generic systems to supply all essential cofactors in addition to the main reactions often requires alteration of several genes that are involved in central metabolic pathways, which often leads to changes that are not beneficial for cell fitness (*14-16*).

To overcome the challenges mentioned above, we proposed a new and minimally perturbing genetic modification approach to enhance several cofactor biosynthesis systems in one go by increasing a pool of glycolytic sugar phosphates that are linked directly to the biosynthesis of NAD(P)H, FAD, FMN, ATP, acetyl-CoA. We proposed that expression of only a single gene encoding a sugar reductase to reduce hexoses to hexitols may result in a rewiring of hexitol metabolism, leading to accumulation of sugar phosphates (Fig. **1a**), precursors of targeted end-products and various cofactors. For sugar supply, we wanted to take advantage of a commonly available sugar, lactose, which is routinely used as an inducer for overexpressing heterologous proteins in *Escherichia coli* (*E. coli*) to supply hexoses (D-glucose and D-galactose). For the choice of sugar reductase, xylose reductase (XR) which is known to reduce several hexoses to generate hexitols was chosen. Although xylose reductase was previously applied in engineered microbes to generate xylitol from xylose and other pentose sugars (*17-19*), its use for increasing levels of a pool of sugar phosphates and for enhancing production of cellular cofactors in engineered cells have never been investigated or proposed.

**Fig. 1.**
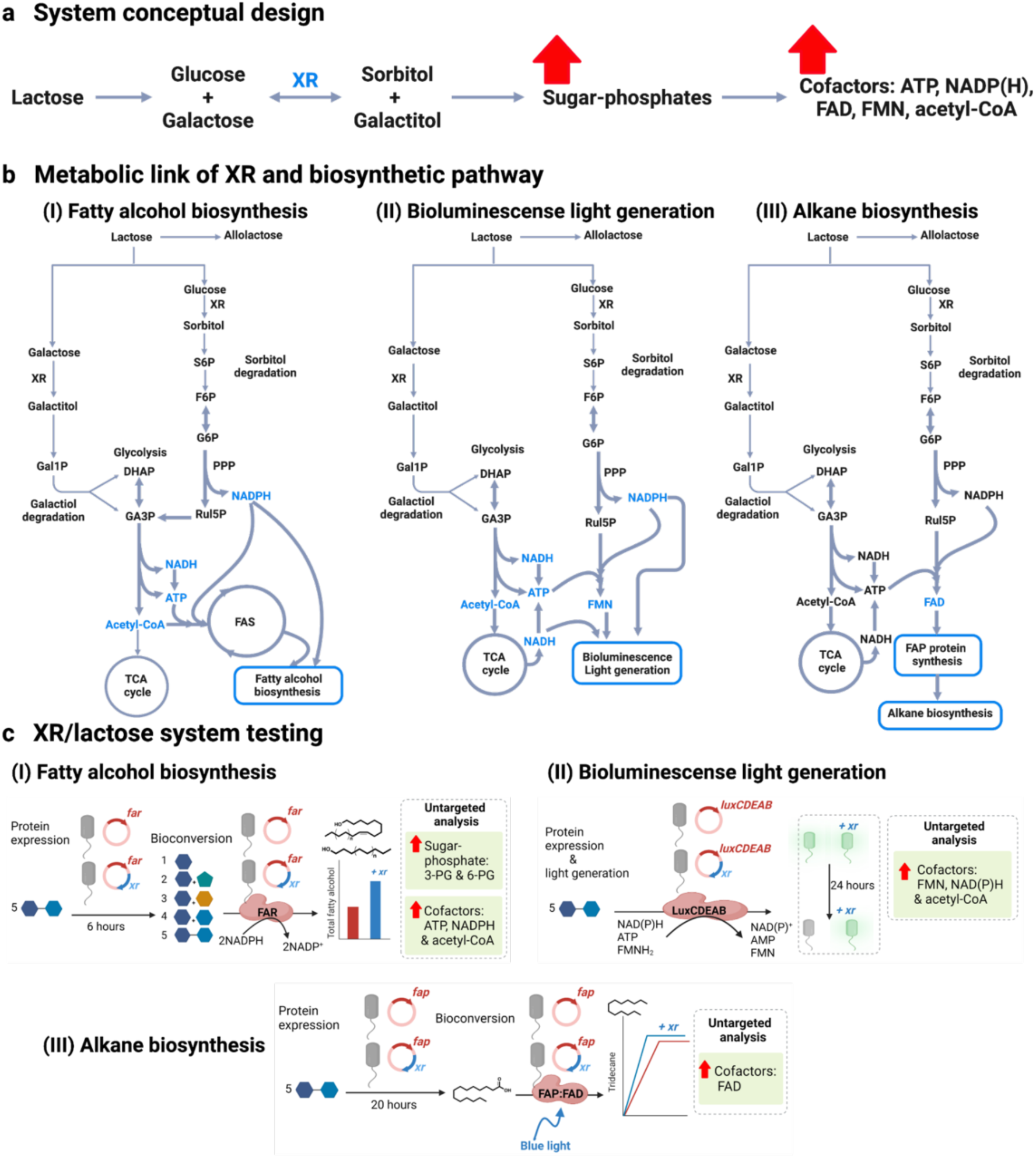
Engineering XR/lactose system for boosting cofactor synthesis for metabolically engineered pathways. **(a)** A conceptual design of using the xylose reductase and lactose (XR/lactose) system to increase sugar-phosphates and thus the synthesis of acetyl-CoA, NAD(P)H/NAD(P), FAD/FMN and ATP cofactors in metabolically engineered cells. **(b)** Links of metabolic and biosynthetic pathways when the XR/lactose is used as a cofactor boosting system for (I) enhancing production of fatty alcohol from the fatty acid biosynthesis having fatty acyl-ACP/CoA as precursors, (II) enhancing light generation by bacterial luciferase using endogenous fatty acid as a substrate, and (III) enhancing the production of alkane using exogenous fatty acid as a substrate. Cofactors that were increased by the XR/lactose boosting system are shown in blue. **(c)** Workflows for testing the functions of the XR/lactose system in enhancing the engineered pathway performance involved measurement of the productivity of fatty alcohol (I), light (II) and alkanes (III), and the changes of whole cell metabolites using untargeted metabolomics. 1, D-glucose; 2, D-glucose/ D-arabinose; 3, D-glucose/ D-fructose; 4, D-glucose/ D-galactose; 5, lactose.

We found that the XR/lactose cofactor boosting system could indeed increase the productivity of three different types of metabolically engineered pathways (fatty alcohol biosynthesis, bioluminescence light generation and alkane biosynthesis) which demand high usage of cofactors such as NAD(P)H, acetyl-CoA, FAD, FMN and ATP in *E. coli* BL21 (DE3) by approximately 2-4-fold. We employed metabolomic and transcriptomic analyses to investigate the metabolic pathways affected by the presence of XR/lactose. The data clearly showed that the XR/lactose boosting system indeed increased levels of intermediates in the sugar phosphate pathways and propagated enhancement effects on key metabolic nodes important for synthesis of common cellular cofactors such as acetyl-CoA, NAD(P)H/NAD(P), FAD/FMN, and ATP. Notably, the patterns of cofactor enhancement are not the same among the three types of cells investigated, but rather customized according to the different demands of the engineered pathways. We further investigated effects of another sugar reductase (glucose dehydrogenase, GDH) in a fatty alcohol production system and found similar enhancement effects as XR, but with less yield enhancement. Altogether, our work demonstrates that the approach of increasing cellular sugar phosphates can be a generic tool to increase *in vivo* cofactor generation to meet cellular demands.

## Results

### Rationale for system design and selection of xylose reductase (XR) and lactose as a versatile cofactor booster

Lactose is a natural sugar routinely used as an inducer for overexpressing heterologous proteins in *Escherichia coli* (*E. coli*) in preparative or industrial scale applications because its cost is markedly cheaper than its analogue IPTG, generally used in lab scale applications (*20-22*). As lactose is commonly added in surplus amount (typically in 2-20 g/L) to maintain protein overexpression, we thus proposed to take advantage of its excess presence as a resource to supply cofactors *via* the reduction of aldose coupled to the synthesis of sugar phosphates and various cofactor biosynthesis in the engineered *E. coli* (Fig. **1a**). When searching for a candidate enzyme to reduce the hydrolyzed products of lactose (D-galactose and D-glucose), XR appeared as an attractive system for performing this task because XR is known to reduce various sugars (*23-25*). We then tested the activity of the purified XR from *Hypocrea jecorina* in the reduction of glucose and galactose using NADPH and found that the enzyme indeed could reduce D-glucose and D-galactose to generate D-sorbitol and D-galactitol (Fig. **S1**). This is consistent with previous work reporting *k*_cat_ values of 4.80 ± 0.20 s^-1^ for D-glucose and 1.28 ± 0.06 s^-1^ for D-galactose reduction (*24*). The ability of XR to reduce the two sugars with *k*_cat_ values within a similar range makes the system a suitable fit for our aims.

With the presence of XR and lactose, cells should be able to generate sorbitol and galactitol at comparable rates; these sugar alcohols would then be converted to sorbitol 6-phosphate (S6P) and galactitol 1-phosphate (Gal1P) *via* D-sorbitol and D-galactitol degradation pathways (hexitol degradation pathways), respectively (Fig. **1b**). We hypothesized that when lactose enters the cell, only a fraction of lactose is converted by β-galactosidase to generate allolactose which binds to the repressor protein and triggers *lac* operon expression. The rest of lactose would be hydrolyzed to D-glucose and D-galactose, which in theory can enter the glycolysis and Leloir pathways, respectively. However, because *E. coli* BL21 (DE3) which widely used as a model organism for studying metabolic pathway engineering cannot utilize galactose as a carbon source due to the *gal* mutation (*26*), the addition of XR provides an additional pathway for the organism to use these hydrolyzed products of lactose efficiently as a resource of cofactors synthesis.

To evaluate the effects of the XR/lactose cofactor boosting system in enhancing the efficiency of biotransformation, we constructed three different *in vivo* biotransformation systems including fatty alcohol synthesis by fatty acyl-CoA reductase (FAR), bioluminescence reporter by bacterial luciferase (LuxCDEAB) and alkane synthesis by fatty acid photodecarboxylase (FAP) (Fig. **1c**). Details for the construction of each engineered pathway are described in the Supplementary Methods. Product yields and the level of cellular metabolite production were measured for these engineered cells in the presence and absence of XR. These systems were chosen because they use different kinds of cofactors. FAR uses two equivalents of NADPH to convert fatty acyl-ACP/CoA to alcohol. LuxAB and LuxCDE generate light upon consumption of FMNH_2_, NAD(P)H and ATP. Different from the other two systems, FAP merely uses FAD as a cofactor and requires external light input to decarboxylate the fatty acid to generate alkane.

### Incorporation of the XR/lactose cofactor boosting system increases the productivity of fatty alcohol

Fatty alcohols are important raw chemicals for industries. They are used as co-emulsifiers in cosmetics, fuel, and food and also as starting materials for various reagents (*27*). We first tested the ability of the XR/lactose system to boost fatty alcohol production in engineered *E. coli* containing the FAR system (*E. coli-far* and *E. coli-far-xr*). After adding lactose to induce protein overexpression for 6 hours, *E. coli*-*far* and *E. coli*-*far-xr* were harvested and employed as biocatalysts for fatty alcohol production (Supplementary Methods). A metabolic pathway of fatty alcohol production of FAR system is shown in Fig. **2a**.

**Fig. 2.**
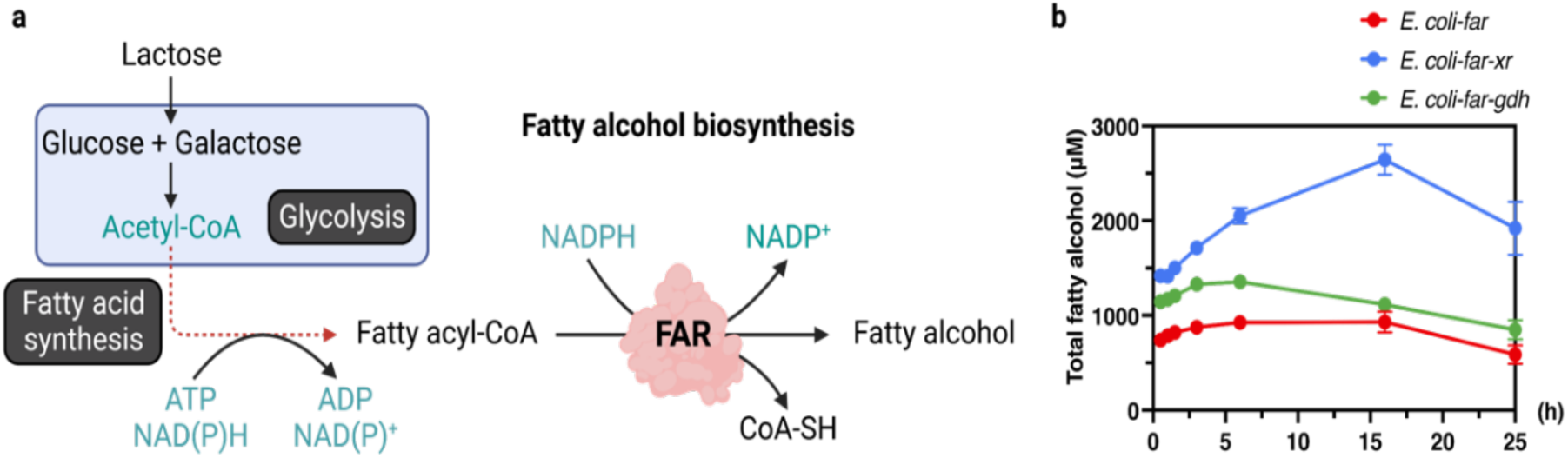
Fatty alcohol production from the FAR-engineering *E. coli*. **(a)** Pathway of fatty alcohol biosynthesis from lactose. Lactose was used as a carbon source for cell growth. Fatty acyl-ACP, an intermediate in fatty acid synthesis, was generated by FAR consumption of NADPH to yield fatty alcohol. **(b)** Production of total fatty alcohol from 10 mM lactose by *E. coli*-*far* (red), *E. coli*-*far-xr* (blue), and *E. coli*-*far-gdh* (green). Data are shown as mean ± s.d., n = 3 replicate cultures.

We first evaluated the impact of sugar carbon sources such as lactose, D-glucose, D-glucose/D-fructose, D-glucose/D-galactose, and D-glucose/D-arabinose on fatty alcohol formation. The results show that *E. coli*-*far-xr* generated more fatty alcohol than *E. coli*-*far* with all of the sugars tested (Fig. **S2**). In the scenario where lactose was used for both protein induction and bioconversion, *E. coli*-*far-xr* generated three times greater fatty alcohol than *E. coli*-*far* (Fig. **2b**). Lactose supplementation at both the protein induction and bioconversion phases resulted in a productivity rate of 165.3 µmol/L/h for *E. coli*-*far-xr* as compared to 58.1 µmol/L/h for *E. coli*-*far* (Fig. **2b**). The total fatty alcohol titer by this engineered cell using lactose was 0.66 mg/ml or 0.18 g/g glucose equivalent in a batch process.

### The XR/lactose system increases the level of sugar-phosphates which enhanced production of acetyl-CoA, ATP, and NADPH required for the fatty alcohol biosynthesis

As the fatty alcohol production by FAR requires two equivalents of NADPH (four electrons reduction process) to reduce fatty acyl-ACP/CoA, an intermediate of the fatty acid synthesis pathway, we hypothesized that the increase of fatty alcohol production might be due to the increase in fatty acyl-ACP/CoA, precursors of fatty acid biosynthesis and cofactors involved in the fatty acid biosynthesis process including NADPH, ATP, and acetyl-CoA (Fig. **2a**). We then performed untargeted metabolomics analysis of *E. coli*-*far-xr* and *E. coli*-*far* cells to compare the differences in their metabolites after 1.5 h of lactose bioconversion to fatty alcohol. These data would present snapshots at early time points of cell metabolite levels and the activities of enzyme networks involved in the fatty alcohol biosynthesis process. Differences observed between *E. coli*-*far-xr* and *E. coli*-*far* cells would indicate changes due to aldose reduction and verify whether the addition of XR could enhance the biosynthesis of sugar phosphates and the biosynthesis of related cofactors.

We performed untargeted metabolomic analysis to identify compounds for which their abundance was different among the two strains. Principal component analysis (PCA) was employed to initially visualize metabolomic data sets. The data clearly showed that the presence of XR indeed caused global metabolic changes in *E. coli*-*far* (Fig. **3a**). Using unpaired and univariate analyses, we found 234 *m/z* features from a total of 1,250 *m/z* features whose fold-changes (FC) were larger than 1.5 (*P* < 0.05) (Fig. **3b**), and 90 out of 234 features could be annotated. The annotated metabolites that displayed significantly different fold-changes were indeed connected to metabolic pathways of sorbitol, galactitol, glycolysis, PPP, phosphopantothenate, CoA biosynthesis and fatty acid syntheses. In addition, we also found an increase of compounds involved with trehalose biosynthesis and glutathione production (Table **S1**).

**Fig. 3.**
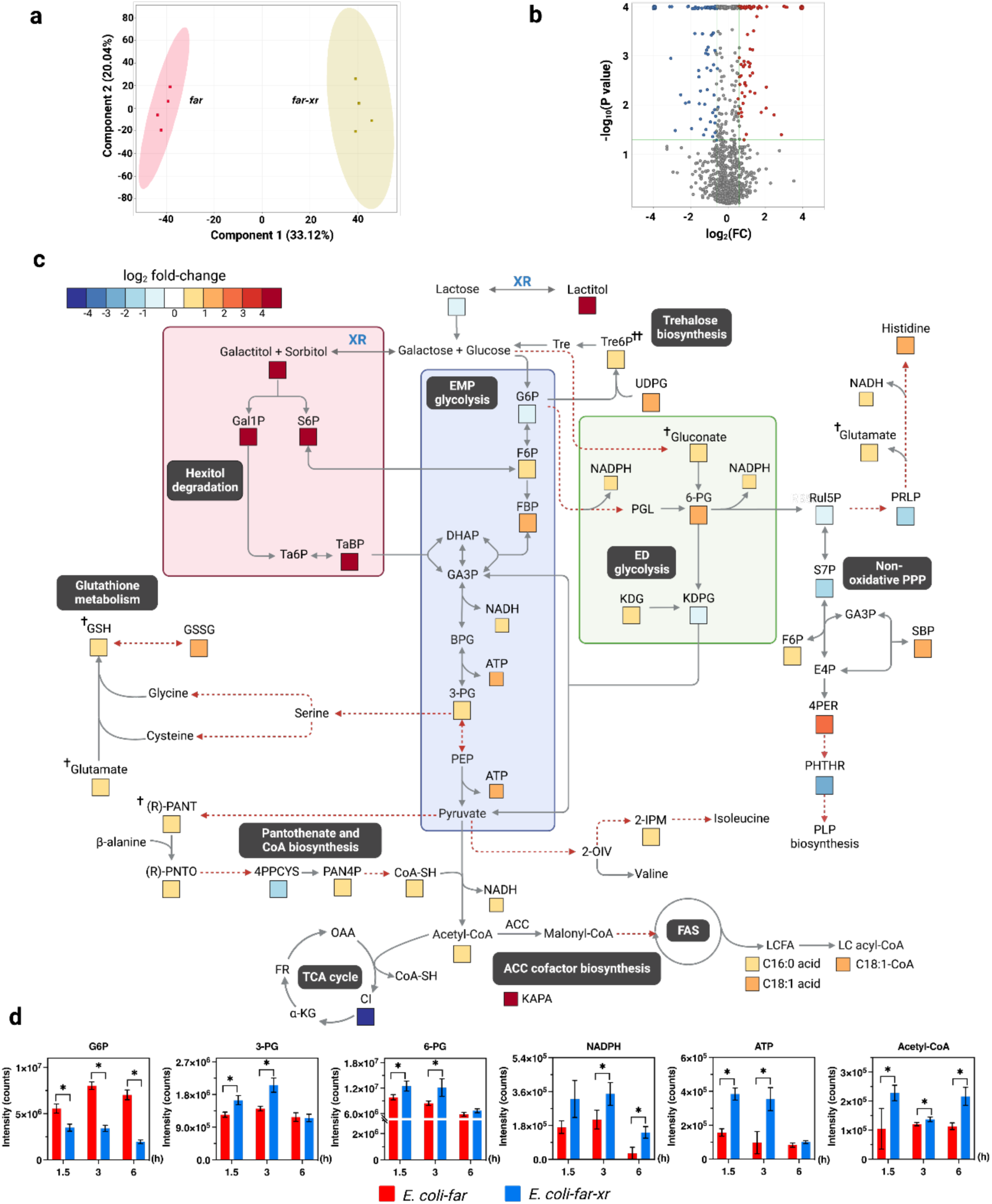
Untargeted metabolomic analysis of the engineered cells for fatty alcohol production with and without XR cofactor enhancement. **(a)** Principal component analysis (PCA) score plots of the first two components show distinct separation of metabolites produced by *E. coli*-*far* and *E. coli*-*far-xr*. Data represent averages from four replicate cultures. **(b)** A volcano plot of metabolites of *E. coli*-*far-xr versus E. coli*-*far* after 1.5 h of the bioconversion process. Red represents the up-regulated metabolites, blue represents the down-regulated metabolites of *E. coli*-*far-xr* compared to those of *E. coli*-*far* (FC > 1.5, *P* < 0.05). Gray represents the metabolites for which the levels are not different between *E. coli*-*far* and *E. coli*-*far-xr*. **(c)** Map of the metabolite changes at 1.5 h after the start of fatty alcohol bioconversion. The log_2_ FC for each metabolite in *E. coli*-*far-xr* compared to *E. coli*-*far* whose FC > 1.5 (*P* < 0.05) are represented according to the color scale. † Denoting metabolites for which FC > 1.2 (*P* < 0.05); †† denoting metabolites for which FC > 1.2 (*P* < 0.1). See all abbreviations in Table **S6. (d)** Time-course analysis of metabolites during the bioconversion process. Data are shown as mean ± s.d., n = 4 replicate cultures; asterisks denote significant differences by multiple *t* test (*P* < 0.05).

To fully understand the changes of metabolites across the pathways involved, we plotted fold-changes (log_2_ FC) of certain metabolites and presented the data based on the magnitude of the log_2_ FC (Fig. **3c**). The time-course data showed that at the beginning of the bioconversion, the levels of acetyl-CoA, ATP, and NADPH that are necessary for the synthesis of fatty acid and fatty alcohol were clearly increased in *E. coli*-*far-xr* as compared to *E. coli*-*far*, with a FC > 1.5 (*P* < 0.05) (Fig. **3c**). Although the amount of these metabolites reduced as the bioconversion progressed, their levels were still greater than those of *E. coli*-*far* (Fig. **3d**). These data clearly suggest that the *E. coli*-*far-xr* reduced glucose and galactose to form sorbitol and galactitol which were then phosphorylated and entered the central carbon metabolisms in the forms of sugar phosphates, S6P and Gal1P, respectively. As shown in Fig. **3c**, the increase of S6P, Gal1P and tagatose 1,6-bisphosphate (TaBP) into glycolysis can directly elevate the level of the glycolytic metabolites, 3-phosphoglycerate (3-PG), and the pentose-phosphate pathway (PPP) metabolites such as 6-phosphogluconic acid (6-PG) (Fig. **3c**), which would allow greater production of acetyl-CoA, ATP, and NADPH to support fatty acid and fatty alcohol synthesis by the FAR system (Fig. **2**). Furthermore, time-course analysis of metabolites related to fatty alcohol synthesis at 1.5 h, 3 h, and 6 h revealed that *E. coli*-*far-xr* generated substantially more (R)-pantoate, 3-PG, and 6-PG than *E. coli*-*far* (Fig. **3d**), indicating that *E. coli*-*far-xr* has greater precursors for synthesis of CoA, acetyl-CoA, and energy related metabolites of cofactors than *E. coli*-*far*.

Interestingly, we also detected the up-regulation of glutathione (GSH) and trehalose 6-phosphate (Tre6P), an intermediate of trehalose biosynthesis, in *E. coli*-*far-xr* (Fig. **3c**). Trehalose and GSH are known to facilitate bacterial adaptability to tolerate oxidative stress (*28*) and osmotic stress (*29-31*), as the overexpression of GSH-encoding genes in *Clostidium acetobutylicum* was shown to improve the strain tolerance to solvent and increase 1-butanol production (*32*). We thus explored tolerance of *E. coli*-*far* and *E. coli*-*far-xr* to H_2_O_2_ and found that *E. coli far-xr* showed higher cell viability than *E. coli*-*far* in the presence of 5 mM H_2_O_2_ (Fig. **S3**). Therefore, the *E. coli*-*far-xr* system which has a greater carbon flow into central metabolism pathways than *E. coli*-*far* without XR can increase the synthesis of GSH and trehalose, which gives it an extra advantage to improve cellular resistance to oxidative and osmotic stressors during the fatty alcohol bioconversion process.

### Enhancement of fatty alcohol production by glucose dehydrogenase (GDH) in comparison to XR

To investigate whether another sugar reductase such as glucose dehydrogenase (GDH) can also be used as a cofactor enhancing system as XR, we constructed the system of *E. coli*-*far-gdh* in which GDH was used as a sugar reductase and investigated fatty alcohol production in comparison to the *E. coli*-*far-xr* system (Supplementary Methods). Both lactose and D-glucose were tested as carbon sources for sugar alcohol production by GDH. The results showed that GDH could also increase the overall fatty alcohol production (∼1.2 fold in the presence of lactose (Fig. **2b**) and about 2-fold in the presence of D-glucose (Fig. **S4**). However, the enhancement effects in fatty alcohol production by the *E. coli*-*far-gdh*/lactose and *E. coli*-*far-gdh*/ D-glucose systems were significantly less than those of the XR/lactose system which was about 3-fold increment (Fig. **S4**).

We then investigated the cause of the difference in product enhancement by XR and GDH by measuring the concentrations of extracellular sugars, intracellular sugars and sugar alcohols resulting from reduction by these sugar reductases (Fig. **S5**). Results (Fig. **S5**) clearly showed that lactose can be up taken by all the cell types tested quite well, while D-glucose was mostly accumulated extracellularly. This might be due to the fact that upon entering the cells, intracellular D-glucose can be quickly metabolized by other metabolic pathways such as glycolysis, rather than being reduced to form D-sorbitol. Interestingly, only the *E. coli*-*far-xr* cells could generate significant amount of D-sorbitol (reduction product of D-glucose) and D-galactitol (reduction product of D-galactose) while the cells containing GDH and D-glucose showed much lower levels of D-sorbitol (Fig. **S5**). A possible explanation is that lactose can be slowly hydrolyzed to generate D-glucose and D-galactose and in the presence of XR, these sugars can be reduced at comparable rates (Fig. **S5**) to generate D-sorbitol and D-galactitol. We noted that in the presence of XR, there was markedly less accumulation of D-galactose in both extracellular and intracellular than in cell types without XR (Fig. **S5**). The results suggest that XR converts D-galactose into D-galactitol which can enter glycolysis *via* DHAP and GA3P, leading to accumulation of sugar phosphates intermediates. All data clearly confirmed the role of XR/lactose as a suitable system for production of sugar alcohols which can lead to enhancement of sugar phosphates and production of cofactors.

### Transcriptomics of genes related to sugar catabolisms of *E. coli*-*far-xr*

All results shown above clearly indicate that the XR/lactose system could increase levels of sugar alcohol and sugar phosphates, resulting in the enhancement of production of various cofactors (Fig. **3**). We then further investigated expression levels (transcriptomics analysis) of genes involving in the early phase of sugar catabolisms where metabolites were shown to be increased in the *E. coli*-*far-xr* cell (Fig. **3**). Cells at the time point of 60 min which should be correlated with metabolites profiles at the time point of 90 min shown in Fig. **3c** were collected for analysis.

Results in Fig. **4** showed that the genes related to gluconate metabolisms (linked to Entner-Doudoroff (ED) and pentose phosphate pathways (PPP) in *E. coli*-*far-xr* were clearly upregulated such as the glucose-6-phosphate dehydrogenase (*zwf*) and the gluconate kinase (*gnt*K) genes which are related to the synthesis of 6-PG (Fig. **4**). The up regulated level of *gnt*K is probably linked to the active PPP pathway in *E. coli*-*far-xr* in which the flux *via* ribulose-5-phosphate (Rul5P) was drawn due to the up-regulation of histidinol-phosphate aminotransferase (*his*C) (Fig. **4**). The up regulation of *his*C would increase the synthesis of histidinol, a precursor of histidine synthesis; this result agreed well with the increased level of histidine shown in Fig. **3**. Similarly, the transcriptomic results can also explain the increased levels of sedoheptulose 1,7-bisphosphate (SBP) and *O*-Phospho-4-hydroxy-L-threonine (4PER) (Fig. **3**) which originated from precursors in the PPP pathway.

**Fig. 4.**
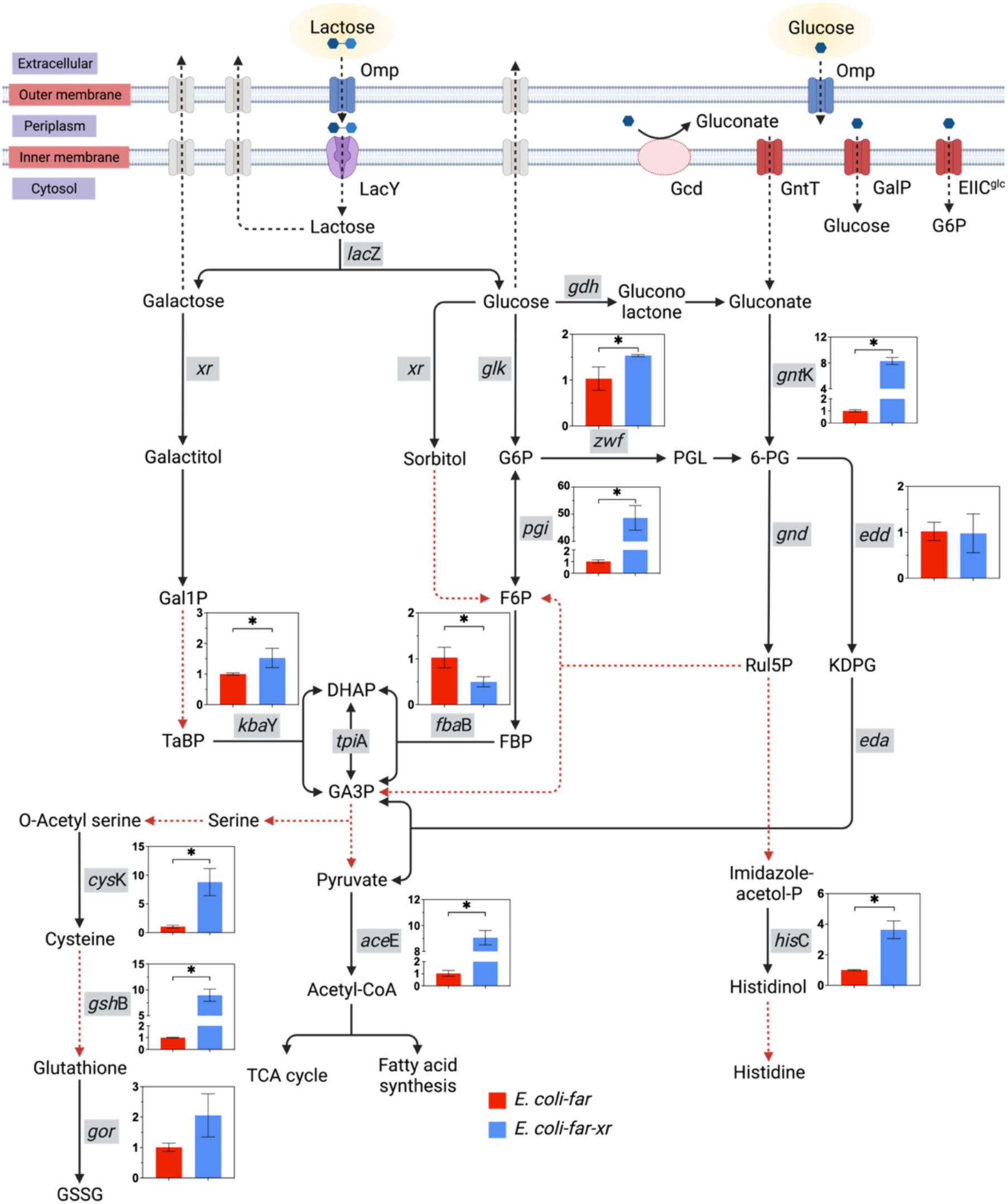
Transcriptomics of sugar catabolism in XR-incorporated cells. Expression levels of genes related to lactose and glucose catabolism in *E. coli*-*far* and *E. coli*-*far-xr* are represented by red and blue bars, respectively. Y-axis of each bar graph refers to the relative quantification (RQ). Genes encoding for enzymes catalyzing relevant steps are labelled in rectangular grey boxes. Black dashed lines indicate sugar metabolic routes. Red dashed lines represent pathways in which multiple enzymatic cascades are possible. All abbreviations are explained in Table **S6**. Data are shown as mean ± s.d., n = 3 replicate cultures; asterisks denote significant differences by multiple *t* test (*P* < 0.05).

Remarkably, the acetyl-CoA synthase gene (*ace*E) was up-regulated 9-fold, explaining the increased levels of acetyl-CoA (Fig. **3d**). The genes involving in synthesis of cysteine and glutathione (*cys*K, *gsh*B, *gor*) were also greatly increased (2-8 folds), in agreement with the increased levels of glutathione observed (Fig. **3c**, **4**). Altogether, the data from the transcriptomic analysis are in agreement with the metabolomic data and could explain the links between the increased levels of D-sorbitol, D-galactitol and sugar phosphates to various pathways leading to synthesis of key metabolites in *E. coli*.

### Incorporation of the XR/lactose cofactor boosting system enhances light generation by bacterial luciferase

To further test the generality of XR/lactose in enhancing cofactors synthesis in other metabolic engineering pathways, we investigated whether XR/lactose could enhance *in vivo* bacterial bioluminescence catalyzed by three enzymatic reactions including those of bacterial luciferase (LuxAB), acid reductase (LuxCDE) and flavin reductase (Fig. **5a**). Flavin reductase generates reduced FMN (FMNH^-^) by reducing FMN using NAD(P)H, while LuxCDE reduces a long chain fatty acid using NADPH and ATP to generate an aldehyde substrate. LuxAB oxidizes a long chain aldehyde to generate acid with concomitant light emission (*33*). The complete catalytic cascade of LuxAB/LuxCDE/flavin reductase has been used for constructing bioreporter systems which are useful for environmental, food and biomedical assessment e.g., detection of toxicant contamination in soil or antibiotic contamination in animal-based food sources. Because this bioreporter system continuously consumes NADPH and ATP to generate light, it provides a good opportunity for testing the ability of the XR/lactose to boost up and supply cofactors to enhance light production (Supplementary Methods).

**Fig. 5.**
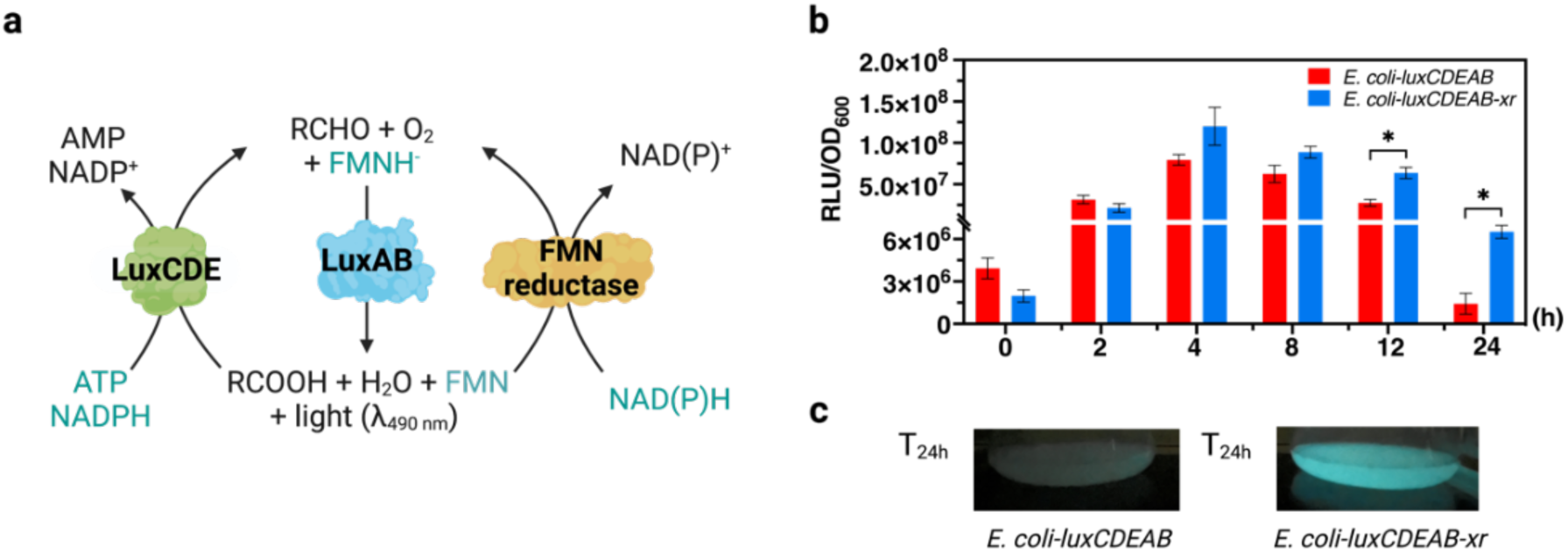
Bioluminescence generation by the bacterial luciferase system. **(a)** The catalytic cycle of complete bacterial luciferase (LuxAB) /acid reductase (LuxCDE)/flavin reductase cascades. **(b)** The bioluminescence generated from *E. coli*-*luxCDEAB* and *E. coli*-*luxCDEAB-xr* in LB media containing 10 mM lactose. Data are shown as mean ± s.d., n = 3 replicate cultures; asterisks denote significant differences by multiple *t* test (*P* < 0.05). **(c)** Culture solutions of the luminous *E. coli*-*luxCDEAB* (left) and *E. coli*-*luxCDEAB-xr* (right) grown at 25°C for 24 hours in LB media supplemented with 10 mM lactose. The photo was taken in the dark.

The results clearly indicate that bioluminescence generated from *E. coli*-*luxCDEAB-xr* is indeed brighter and lasts longer than that from *E. coli*-*luxCDEAB* (Fig. **5a**). Although, both cell types achieved their highest bioluminescence levels after 4 h, the brightness of *E. coli*-*luxCDEAB-xr* was 1.5-fold higher than of *E. coli*-*luxCDEAB*, and the light signal of *E. coli*-*luxCDEAB-xr* remained after 24 h, whereas that of *E. coli*-*luxCDEAB* was almost undetectable (> 4-fold enhancement) (Fig. **5b**). This is due to the ability of the XR/lactose system to generate more precursors for the biosynthesis of ATP, NAD(P)H, and FMNH^-^ thus better prolonging light generation in the *E. coli*-*luxCDEAB-xr* (see details of metabolomics analysis in the section below).

### The XR/lactose cofactor boosting system enhances light production *via* increased FMN, ATP and NAD(P)H

To understand the molecular mechanisms of bioluminescence enhancement by the XR/lactose boosting system, we performed untargeted metabolomic analysis to compare the difference in metabolite levels between *E. coli*-*luxCDEAB* and *E. coli*-*luxCDEAB-xr*. For a time point of investigation, we chose to investigate the cells after 24 h of light generation because it was a period where *E. coli*-*luxCDEAB-xr* showed the greatest difference in light signals. Details of methodology are described in Supplementary Methods and the overall analysis protocols were similar to those previously described for fatty alcohol production.

The results of PCA analysis of metabolites derived from *E. coli*-*luxCDEAB* and *E. coli*-*luxCDEAB-xr* can be clearly grouped according to each cell type (Fig. **6a**). Using unpaired and univariate analyses, we found 353 *m/z* features from a total of 1,269 *m/z* features whose FC > 1.5 (*P* < 0.05) (Fig. **6b**), and 93 of the 353 *m/z* features could be annotated. Consistently, metabolites altered were mostly similar to those pathways observed in fatty alcohol production shown in Fig. **3** (Fig. **6c**, Table **S2**). Interestingly, we also found additional altered pathways in bioreporter cells including biosynthesis of FMN, folate biosynthesis, purine and pyrimidine, and peptidoglycan (Fig. **6c**, Table **S2**).

**Fig. 6.**
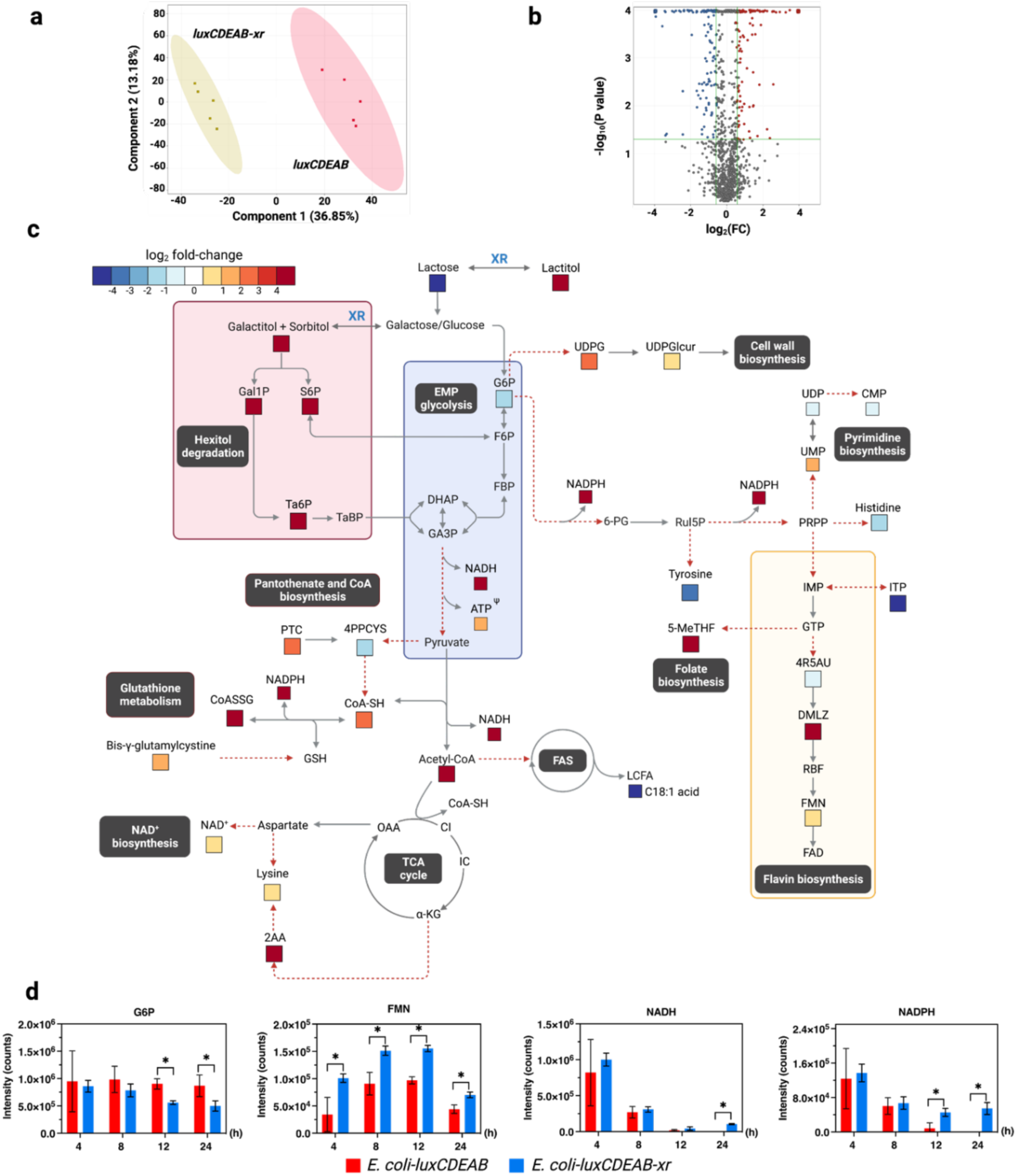
Metabolomic analysis of bioluminescence bioreporter cells. **(a)** Principal component analysis (PCA) score plots of metabolite levels of *E. coli*-*luxCDEAB* and *E. coli*-*luxCDEAB-xr* after 24 h of bioluminescence generation. PCA of the first two components shows distinct separation of variances which indicate the influence of the types of metabolites produced by *E. coli*-*luxCDEAB* and *E. coli*-*luxCDEAB-xr*. Data are shown as mean ± s.d., n = 5 replicate cultures. **(b)** Volcano plots of metabolites in *E. coli*-*luxCDEAB* and *E. coli*-*luxCDEAB-xr* cells at 24 h after bioluminescence generation. Red represents the up-regulated metabolites, while blue represents the down-regulated metabolites as compared to *E. coli*-*luxCDEAB* (FC > 1.5, *P* < 0.05). Gray represents metabolites which are not different between *E. coli*-*luxCDEAB* and *E. coli*-*luxCDEAB-xr*. **(c)** Map of the metabolic changes in bioluminescent cells after adding lactose for 24 h. The log_2_ FC for each metabolite in *E. coli*-*luxCDEAB-xr versus E. coli*-*luxCDEAB* whose FC > 1.5 (*P* < 0.05) are displayed according to the color scale. ^Ψ^ Denoting the level of metabolites derived from LC/MS QQQ analysis. A full list of abbreviations is described in Table **S6. (d)** Time-course analysis of metabolites after adding 10 mM lactose. Data are shown as mean ± s.d., n = 5 replicate cultures; the asterisks denote significant differences as determined by multiple t test (P < 0.05).

Comparison of levels of intracellular metabolites of the two prototype cells using the fold-change analysis showed that *E. coli*-*luxCDEAB-xr* has higher levels of metabolites related to biosynthesis pathways for production of NADPH (FC > 1.2, *P* < 0.1) and FMN (FC > 2, *P* < 0.05) than those of *E. coli*-*luxCDEAB* (Figs. **6c**,**d**). Here, we could not observe ATP; this might be due to the rate of ATP utilization being much greater than the rate of production, causing the level of ATP to be lower than the detection limit of the high-resolution mass spectrometer (IM-QTOF, LC-MS system). We thus used triple quadrupole (QQQ) mass spectrometry operating in selective ion monitoring (SIM) mode for targeted analysis and could observe that the level of ATP in *E. coli*-*luxCDEAB-xr* was 2-fold higher than that of *E. coli*-*luxCDEAB* at 12 hours and 24 hours of light generation (Fig. **S6**). The metabolomic data shown in Fig. **6c** also demonstrate that the light enhancement is likely due to the increase of S6P and Gal1P, which are directly linked to the increased levels of NADPH and FMN. The decrease in the level of G6P, a branch metabolite from glycolysis, and PPP was observed, indicating that G6P was heavily consumed by the *E. coli*-*luxCDEAB-xr* for creating NADPH and FMN. The increased presence of these metabolites can be used in the bioluminescence catalytic cascade and thus enhances bioluminescence light generation.

High detected concentrations of metabolites in folate biosynthesis such as 5-methyltetrahydrofolate (5-MeTHF) was in agreement with the high levels of NAD(P)H and FMN, indicating that PPP is highly active in the XR-incorporated biocatalyst which is due to the increase in G6P flow into the PPP biosynthesis (Fig. **6c**). 5-MeTHF is necessary for production of a number of cellular components such as thymidylate, pantothenate, and purine nucleotides. The increased level of 5-MeTHF possibly results in the observed up-regulation of acetyl-CoA, fatty acid, purine and pyrimidine and peptidoglycan biosynthesis in *E. coli*-*luxCDEAB-xr* shown in Fig. **6c** and Table **S2**. These compounds are important for the synthesis of cellular components during growth, thus enhancing prolonged and brighter light production signals.

A time-course analysis of metabolites obtained from *E. coli*-*luxCDEAB-xr* and *E. coli*-*luxCDEAB* cells after addition of lactose for 4, 8, 12, and 24 h revealed the decreased levels of G6P in *E. coli*-*luxCDEAB-xr*, suggesting a fast conversion of this sugar phosphate into the PPP pathway which helps prolong bioluminescence production. The brighter light of *E. coli*-*luxCDEAB-xr* than that of *E. coli*-*luxCDEAB* was also supported by the higher levels of FMN and NADPH in *E. coli*-*luxCDEAB-xr* than those of *E. coli*-*luxCDEAB* throughout 4-24 h (Fig. **6d**). Altogether, these results agree well with the fatty alcohol production system in that the XR/lactose system can boost generation of cofactors required by the engineered metabolic pathway, including NAD(P)H, ATP, and acetyl-CoA.

### Incorporation of XR enhances the alkane production rate

The third system we used for testing cofactor boosting effects of XR/lactose was fatty acid photodecarboxylase (FAP), a flavoenzyme catalyzing photodecarboxylation of fatty acids to produce hydrocarbons. As illustrated in Fig. **7a**, this enzyme requires photons from blue light (400-520 nm) to activate its FAD cofactor for catalysis. Our results showed that the combined use of FAP and XR led to improvement of *in vivo* production of alkane from exogenous fatty acid. We tested two different systems of XR expression, plasmid and genome integrated expression. The results showed that both systems could enhance the production of tridecane (see Fig. **S7** for the plasmid system). For the genome-integrated XR, we found a 1.6-fold improvement of *in vivo* production of tridecane from exogenous tetradecanoic acid by *E. coli*-*fap-xr* compared to *E. coli*-*fap*. The production rate was increased from 0.8 mmol/L/h to 1.3 mmol/L/h (Fig. **7b**) and the period required for completing the reaction by the FAP-XR system was much shorter than that of the system without XR. As FAD is a key cofactor in photodecarboxylation by FAP, we hypothesized that co-expression of XR enhances riboflavin synthesis, which in turn increases the level of FAD-bound FAP.

**Fig. 7.**
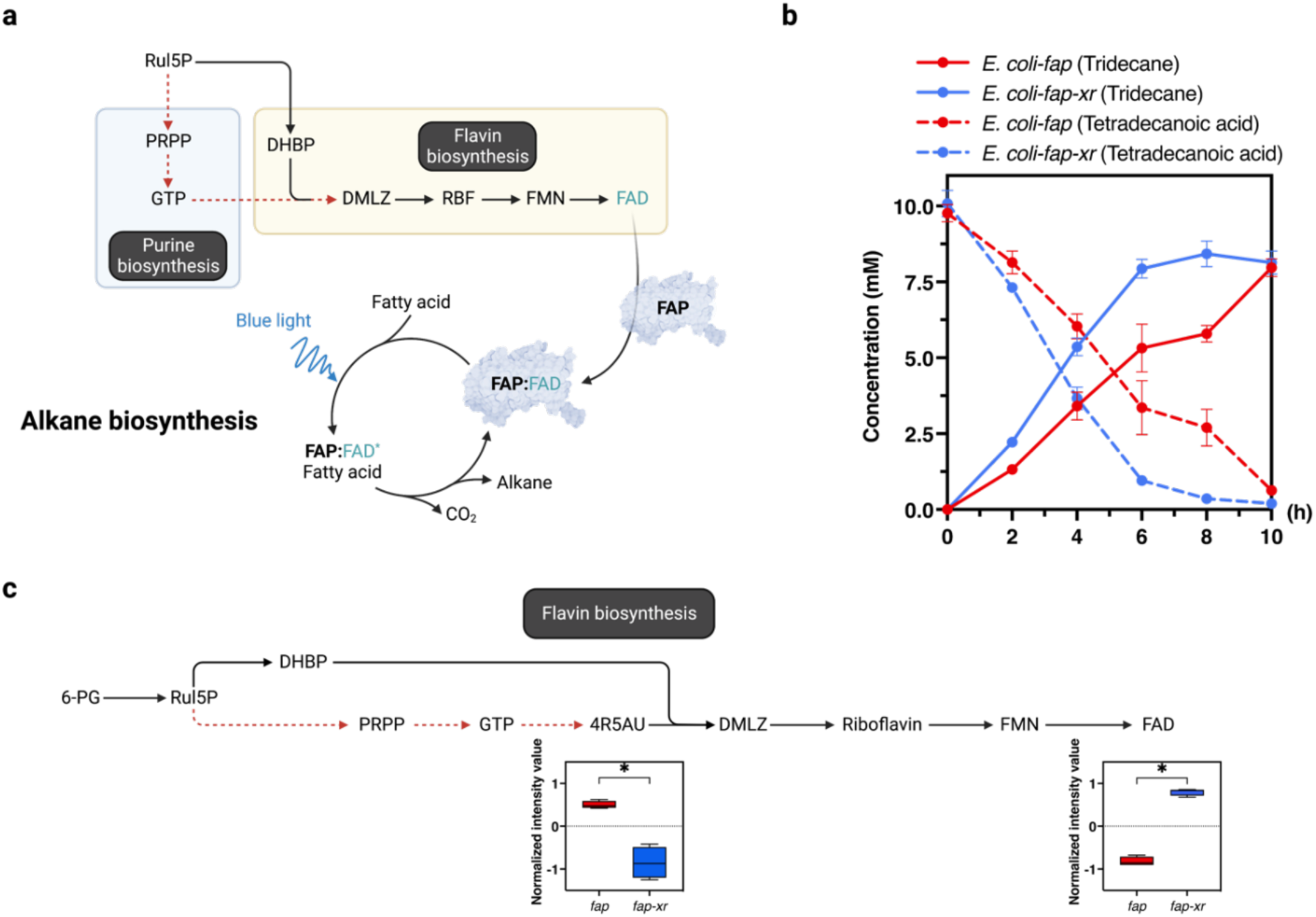
Alkane biosynthesis by fatty acid photodecarboxylase. **(a)** Pathways involved with FAD synthesis and the catalytic cycle of alkane biosynthesis by fatty acid photodecarboxylase (FAP). The reaction requires blue light (450 nm) to activate FAD at the enzyme active site for decarboxylation of fatty acid substrates to form alkanes and CO_2_. **(b)** Production of alkane from fatty acid by *E. coli* harboring *Cvfap*. Tridecane production by *E. coli*-*fap* is indicated by the red-solid line and production by *E. coli*-*fap-xr* is indicated by the blue-solid line. Utilization of tetradecanoic acid by *E. coli*-*fap* is shown as the red-dashed line, and *E. coli*-*fap-xr* as the blue-dashed line. The bioconversion processes were carried out in potassium phosphate buffer containing 10 mM tetradecanoic acid. Data are shown as mean ± s.d., n = 4 replicate cultures. **(c)** Comparison of intracellular flavin-related metabolites after 5 hour of alkane bioconversion. Levels of metabolites were normalized by total abundance from 4 replicates and represented in a box-whisker plot. The asterisks denote significant differences as determined by a multiple *t* test (*P* < 0.05). See abbreviations in Table **S6**.

### XR accelerates alkane production rate by increasing the formation of FAD in *E. coli*

Because the addition of XR increases the rate of alkane production of *E. coli* harboring FAP, we employed untargeted metabolomics analysis to investigate the function of XR in this biocatalyst and during the bioconversion process. We found that XR altered the flavin biosynthesis pathways, resulting in the higher level of FAD in *E. coli*-*fap-xr* compared to that in *E. coli*-*fap* (Fig. **S8**). Analysis of *E. coli*-*fap-xr* and *E. coli*-*fap* cells after 5 h of the bioconversion process, at which point the alkane productivity was increased the most, revealed that the difference between these two cell types indeed lies mostly in the series of metabolites involved in FAD biosynthesis (Table **S3**). The intracellular concentration of 5-amino-6-ribitylamino uracil (4R5AU), an intermediate of flavin biosynthesis, was less accumulated in *E. coli*-*fap-xr* than in *E. coli*-*fap*, and the level of 4R5AU in *E. coli*-*fap-xr* was 2.5-fold lower (*P* < 0.05) than that of *E. coli*-*fap*. However, *E. coli*-*fap-xr* had a level of FAD 3-fold higher (*P* < 0.05) than that of *E. coli*-*fap* (Fig. **7c**). These data implied that the FAD biosynthesis in *E. coli*-*fap-xr* was more active than that of *E. coli-fap*, resulting in higher amounts of the active biocatalyst, FAD-bound FAP, for catalyzing the conversion of fatty acid to alkane. Confirmed by the whole-cell bioconversion data, *E. coli*-*fap-xr* showed a higher rate of bioconversion than *E. coli*-*fap* (Fig. **7b**).

The increased rate of alkane bioconversion by *E. coli*-*fap-xr* allows a shorter light exposure time for the bioconversion process. This would allow the system to gain higher efficiency in alkane production because FAP can be inactivated by radical mechanisms after prolonged light exposure (*34*). In the past, several methods have been attempted to maintain bioconversion by prolonging the substrate incorporation step to increase the FAP stability (*35*) or supplying new active cells to the reactor (*36, 37*). With the XR/lactose boosting system, the alkane production by FAP can be enhanced using a simple method.

## Discussion

The results presented here demonstrated the efficient use of XR/lactose as a simple tool to supply various cofactors in *E. coli* for enhancing product formation by metabolically engineered cells requiring different quantities of cofactors (e.g., acetyl-CoA, NAD(P)H, ATP, and FAD/FMN). Fatty alcohol biosynthesis by fatty acyl-CoA reductase (FAR), light generation by bacterial luciferase (LuxCDEAB), and alkane biosynthesis by fatty acid photodecarboxylase (FAP) were chosen as systems to validate this tool. Our results showed that XR/lactose had a strong capability to drive metabolically engineered cells in two settings: (1) in the multi-step bioconversion, in which engineered cells require a steady supply of cofactors throughout the process and (2) in the single-step bioconversion in which the engineered cells only require a cofactor for producing an active protein. Another interesting aspect of the XR/lactose system is that the changes of metabolites are not the same for all systems; the metabolic flux appeared to change according to the systems’ particular demand.

For the multi-step bioconversion, the data herein showed the effectiveness of XR/lactose in boosting the production of fatty alcohol and light. By using the XR/lactose system, the fatty alcohol-producing cell showed a 3-fold increase in productivity, while the light-generating cell could emit light even after 24 hours, ∼4-fold brighter light than the system without XR/lactose. Levels of acetyl-CoA, NADPH, and ATP were found to be higher in the XR/lactose systems in both fatty alcohol and light-generating cells because both systems require high utilization of these cofactors. Interestingly, only the light-generating cells showed an increase of FMN because this cofactor is required for the function of LuxAB.

For the single-step bioconversion, as for the case of *E. coli* containing FAP, our results revealed that only FAD was found as a cofactor prominently changed in this cell type. The amount of FAD in the cell using XR/lactose was around 3-fold higher than in the control cell. The alkane productivity was improved by 2-fold.

Among the metabolites detected by the untargeted metabolomics approach, S6P/Gal1P and TaBP, the products of sorbitol and galactitol metabolic pathways which can enter glycolysis, existed in all cells with similar time-course kinetics of production. The data suggest that the XR cell can convert D-glucose and D-galactose to the corresponding sugar alcohols efficiently and simultaneously. An increase in sugar monophosphates (e.g., G6P and 3-PG) and PPP monophosphates pools (e.g., 6-PG) was observed as one of the core changes in fatty alcohol and light-producing cells. Moreover, only the fatty alcohol-producing cells using XR/lactose showed a high level of 2-Isopropylmalic acid (an intermediate of valine and leucine biosynthesis, Table **S1**) and histidine. 2-Isopropylmalic acid and histidine are derived from their carbohydrate precursors from the glycolysis and PPP pathways, respectively. The data imply that the changes of metabolites in the XR cell are according to the cell-specific demand. We propose that this is the “user-pool” mechanism in which the activities of central metabolic pathways are altered based on utilization or depletion of particular compounds by the engineered pathway. Without utilization, the XR cell does not increase cofactors universally. Due to their activities in the sugar metabolic pathways, the XR/lactose system also increased cellular levels of metabolites required for alleviating cellular stress such as glutathione and CoA disulfide, thus increasing cell fitness and allowing the XR-incorporated cell to tolerate to H_2_O_2_ better. Therefore, the XR/lactose system is a simple and useful technology for enhancing cell fitness for bioproduction.

The use of the XR/lactose system to enhance cofactor production according to specific cellular demand or user-pool mechanisms is different from previous engineering efforts to increase cellular cofactors which generally involved manipulation of several genes in central metabolic pathways (*7, 38, 39*). Satowa *et al*. (*7*) overexpressed *atoB* (encoding acetyl-CoA acetyltransferase) and disrupted *pgi* gene in order to increase the level of acetyl-CoA and NADPH, respectively. Although the ratio of NADPH/NADP was elevated by 7-fold, the engineered strain does not show an improvement in mevalonate production. Additional deletion of *gltA* (encoding citrate synthase) and overexpression of *atoB* were required to increase the mevalonate titer in *E. coli* around 1.5-fold. In order to increase the pools of precursors e.g., GA3P and pyruvate for lycopene production in *E. coli*., overexpression of nine genes in the central metabolic pathways were required (*38*). The genetic modifications for boosting cofactors production are not always successful due to several disturbance of metabolic pathway largely in central carbon metabolisms which resulting in deteriorate cell growth. Furthermore, functions of those cofactors in metabolic network make genetic engineering entangling.

It should be noted that the use of the XR/lactose system for enhancing the production of cofactors here is different from previous studies of cells harboring XR. Most studies employed XR in combination with xylose (*40-42*) to reduce xylose to xylitol which is a valuable compound. Sun *et al*. (*42*) co-expressed acetyl-CoA synthase (ACS) with XR, xylitol dehydrogenase (XDH), and xylulose kinase (XK) in *Saccharomyces cerevisiae* to enable the co-utilization of xylose and acetate as a carbon source. It was interesting to note that in the previous study in which XR was incorporated into cells harboring the α,w-dicarboxylic acids (DCA) pathway to produce DCA from tetradecanoic acid in *E. coli*, the DCA titer was increased by 1.8-fold. No clear reasons were available then.

Based on this current data, we hypothesize that the effectiveness of XR/lactose in cofactor boosting effects may be due to its ability to mitigate the glucose repression in *E. coli*. Glucose repression generally happens in bacteria to repress the metabolic pathways involved in the utilization of alternative carbon sources (*43*). Analysis of intracellular and extracellular lactose, D-glucose, D-galactose, and D-sorbitol/D-galactitol levels clearly showed that the XR/lactose system can reduce D-glucose/D-galactose to generate D-sorbitol/D-galactitol effectively (Fig. **S5**). Genes related to the catabolisms of galactitol such as *kbaY* (encoding tagatose-1,6-bisphosphate aldolase 1 which catalyzes the reversible aldol condensation of DHAP and GA3P to form TaBP (an intermediate of galactitol metabolism) was upregulated 14-fold in 60 min as compared to at 0 min (*P* < 0.05), respectively (Fig. **S9**). These data also support that the metabolic activities for converting D-sorbitol/D-galactitol into sugar phosphates were enhanced.

In conclusion, our findings suggest that the XR/lactose system can be used as a simple synthetic biology tool to enhance levels of sugar phosphates which lead to *in situ* generation of various cofactors upon cellular demand (Fig.**8**). The synthesis of specific cofactors depends on cellular usage of overexpressed metabolic pathways. As the demand for cellular cofactors is dynamic, our approach of increasing precursor pools of sugar-phosphates would reduce the metabolic burden and eliminate cofactor imbalance in the metabolically engineered cells.

**Fig. 8.**
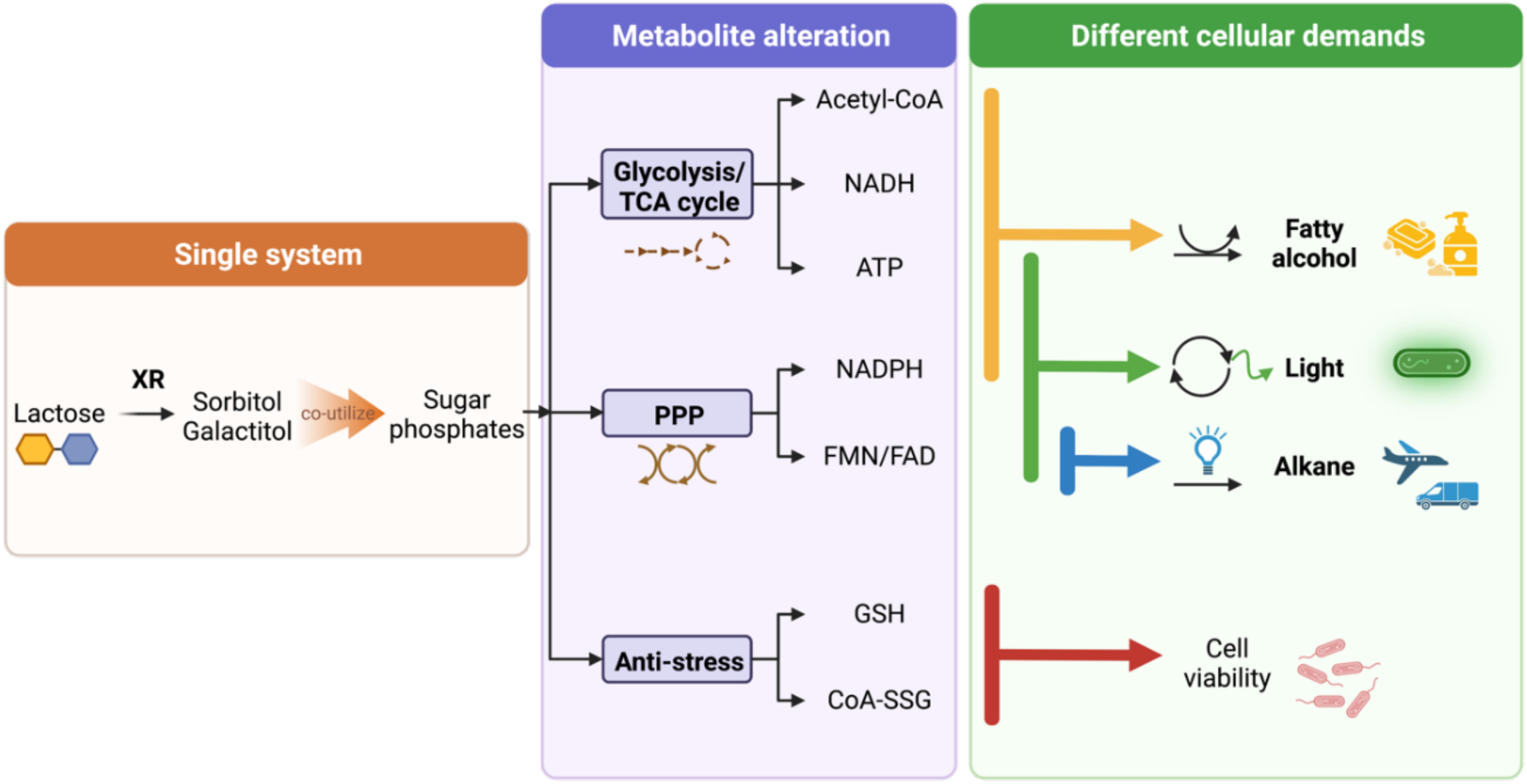
A summary of the XR/lactose system, a simple synthetic biology tool for increasing sugar phosphate levels and producing different cofactors on demand.

## Materials and Methods

### Reagents

All commercially available chemicals were purchased from Agilent, Sigma-Aldrich, TCI, or Merck. T4 DNA ligase, Restriction enzymes, PCR kits, Gibson kits, and other molecular biology reagents were purchased from New England Biolabs (NEB). Plasmid extraction mini kit and Gel/PCR mini kit were purchased from Favogen Biotech Corporations.

### Strains and plasmids

BL21 (DE3) (*fhuA2 [Ion] ompT gal (λ DE3) [dcm] ΔhsdS*) cells from Novagen was used for protein expression. The pET28a, pETDuet-1, pRSFDuet-1, and pCDFDuet-1 vectors (Novagen) were used for construction of expression plasmids. Every gene on the pET28a and Duet-1 vector was controlled by the T7 ribosome binding site (RBS) sequence, AAGGAG. The preparation of expression plasmids used in this study is described below and shown in Fig. **S9**.

### Construction of pETDuet-*mbp-far*, pETDuet-*mbp-far-gst-xr*, and pETDuet-*mbp-far-gdh*

For construction of pETDuet-*mbp-far* plasmid, a gene encoding for a fusion protein of maltose binding protein (MBP)-fatty acyl coA reductase (FAR) from *Marinobacter aquaeolei* VT8 was amplified with the primers F_lacI and F-M13 (Table **S4**) using pMAL-*mbp-far* as a template. The *mbp-far* gene was codon-optimized, synthesized, and supplied on a pMAL plasmid by Genscript. The resulting PCR product was digested with *Nde*I and ligated with the *Nde*I/*Eco*RV cut pETDuet-1 vector to yield the pETDuet-*mbp-far* plasmid. For construction of the pETDuet-*mbp-far-gst-xr* plasmid, the pETDuet-*mbp-far* was digested with *Not*I/*Nco*I and ligated with the PCR product of *gst-xr* genes using the Gibson assembly kit. The *gst-xr* from *Hypocrea jecorina* gene was codon-optimized, synthesized, and supplied on a pGEX-4T plasmid by Genscript. The primers F-gst-xr and R-gst-xr (Table **S4**) were used to amplify the *gst-xr* gene using the pGEX-*gst-xr* as a template. The glucose dehydrogenase (*gdh*) gene from *Bacillus amyloliquefaciens* SB5 was previously constructed in the pJET1.2/blunt-end cloning vector (*1*). The primers F-ATG-6XHis-gdh and R-ATG-6XHis-gdh (Table **S4**) were used to amplify the *gdh* gene and subcloned into pRSFDuet-1 using the Gibson assembly kit at *Nco*I and *Not*I sites to yield the pRSFDuet-6xHis-*gdh* plasmid. For construction of pETDuet-*mbp-far-gdh*, pRSFDuet-6xHis-*gdh* was digested and the resulting fragment was ligated into pETDuet-*mbp-far* at *Apa*I and *Nde*I cutting sites.

### Construction of pETDuet-*luxCDEAB*

For construction of the pETDuet-*luxCDEAB* plasmid, genes encoding for *LuxCDE* and *LuxAB* from *Vibrio campbellii* (*Vc*) *were* amplified with the primers F-luxCDE and R-luxCDE and primers F-luxAB and R-luxAB, respectively (Table **S4**). pET17b-*VcluxCDE* and pET11a-*VcluxAB* plasmids were used as templates for amplification. The resulting PCR product of *luxAB* was digested with *Nde*I/*Xho*I and ligated with the *Nde*I/*Xho*I digested pETDuet-1 vector to yield the pETDuet-*luxAB* plasmid. The resulting pETDuet-*luxAB* was cut with *Bam*HI/*Nco*I and ligated with *luxCDE* PCR product using Gibson assembly kit to yield the pETDuet-*luxCDEAB* plasmid.

### Construction of pRSFDuet-*gst*-*xr*

For construction of the pRSFDuet-*gst*-*xr* plasmid, a codon optimized *gst-xr* from *Hypocrea jecorina* was amplified with the primers F-pRSF-xr and R-pRSF-xr (Table **S4**) using pGEX-*gst-xr* as a template. The resulting PCR product was ligated with *Nde*I digested pRSFDuet-1 vector using a Gibson assembly kit to yield the pRSFDuet-*gst-xr* plasmid.

### Construction of pET28a-*fap* and pCDFDuet-*fap*

pET28a-*fap* plasmid was prepared according to previous reports(*2, 3*). Briefly, cDNA encoding FAP from *Chlorella variabilis* (*Cvfap*) was cloned, amplified by excluding predicted targeting sequence for chloroplasts, and codon-optimized for expression in *E. coli* and ligated into the pLIC07 vector. The gene cassette consists of 6xHis-tagged, thioredoxin, a tobacco etch virus (TEV) protease cleavage site and a gene encoding for *Cvfap*. This construct was cloned into pET28a between *Nde*I/*Hin*dIII sites to generate pET28a-*fap*. For construction of the pCDFDuet-*fap* plasmid, pET28a-*fap* was cut with *Nde*I and *Xho*I to generate a fragment consisting of 6xHis-tagged, thioredoxin, TEV protease cleavage site, and *Cvfap*. This fragment was inserted into the second multiple cloning site (MSC-2) of pCDFDuet-1 between *Nde*I/*Xho*I sites by T4 DNA ligase.

### Construction of *E. coli* BL21 (DE3), *glms::*pT7-*gst*-*xr, Gm*^*r*^

To construct *E. coli* BL21 (DE3), *glms::*pT7*-gst-xr, Gm*^*r*^ strain, the pT7*-gst-xr* fragment was amplified with the primers F-pT7-gst-xr and R-pT7-gst-xr (Table **S4**) using pETDuet-*mbp-far-gst-xr* as a template. This PCR fragment was ligated with *Sma*I cut pUC18T-mini-*Tn7T-Gm*^*r*^*-R6K*, generating pUC18T-mini-*Tn7T-Gm*^*r*^*-R6K-pT7-gst-xr*. The resulting pUC18T-mini-*Tn7T-Gm*^*r*^*-R6K-pT7-gst-xr* plasmid was used to transform *E. coli* BW20767. *E. coli* BW20767 containing pTNS2 and *E. coli* BW20767 containing pUC18T-mini-*Tn7T-Gm*^*r*^*-R6K-pT7-gst-xr* was used to transfer pT7-*gst*-*xr* into *E. coli* BL21 (DE3) containing the pACYCDuet-1 plasmid by triparental conjugation. *E. coli* transconjugants harboring chromosomal insert of the *pT7-gst-xr* gene at downstream of the *glmS* genes were selected on LB agar plate containing 30 μg/ml gentamycin while *E. coli* BW20767 was counter-selected using 15 μg/ml chloramphenicol. The *E. coli* BL21 (DE3), *glms::pT7-gst-xr, Gm*^*r*^ strain was subsequently streaked on an LB agar plate containing 30 μg/ml gentamycin for 10 passages for removing the pACYCDuet-1 plasmid. Colony PCR was used to identify clones with the insertion of *pT7-gst-xr* on the chromosome.

### Preparation of fatty alcohol producing cells and bioconversion assays

The pETDuet-*mbp-far* plasmid was transformed into competent *E. coli* BL21 (DE3) with 15 ng of plasmid to generate the engineered cell without *xr*. The pETDuet-*mbp-far-gst-xr* plasmid was transformed into competent *E. coli* BL21 (DE3) with 15 ng of plasmid to generate the engineered cell with *xr*. The pETDuet-*mbp-far-gdh* plasmid was transformed into competent *E. coli* BL21 (DE3) with 15 ng of plasmid to generate the engineered cell with *gdh*. The transformed cells were selected by growth on LB with 100 µg/ml ampicillin. A single colony was selected and grown in LB with 100 µg/ml ampicillin as a starting liquid culture overnight at 37 °C. The corresponding variant was then sub-cultured into TB media with 1% inoculant (by volume) and grown at 37 °C until the OD_600_ reached 0.6. The culture was then induced with 1 mM lactose and incubated further at 25 °C for 6 h. Bacterial cells were harvested by centrifugation at 4 °C, and then resuspended in 0.1 M potassium phosphate (Kpi) buffer pH 7.5 to obtain a cell suspension with OD_600_ of 30 for 1 mL. This suspension was used for biocatalytic reactions. Various sugars including 20 mM glucose, 10 mM glucose/10 mM arabinose, 10 mM glucose/10 mM fructose, 10 mM glucose/10 mM galactose, and 10 mM lactose were used as carbon sources. The reactions were incubated at 220 rpm, 25°C. The biocatalytic reaction (1 mL) was carried out in a gas tight vial for designated periods of time and quenched by adding 2 mL ethyl acetate solvent containing internal standards (200 µM tetradecane) with vigorous mixing. Upon addition of ethyl acetate, the mixture was centrifuged (4 °C, 3900g, 20 min) to enhance phase separation and the upper organic layer was analyzed by Agilent7890B GC-MS equipped with a HP5-MS column. The samples for sugar and gene expression analysis (100 µL) were collected from the reaction and centrifuged at 4000g at 4°C for 10 min. For intra- and extracellular sugar analysis, cell pellet and supernatant were separated and kept at -20°C. For gene expression analysis, only cell pellet was snap-frozen in liquid nitrogen and kept at -80°C until further analysis.

### Light generating cell preparation and light measurement

The pETDuet-*luxCDEAB* plasmid was transformed into competent *E. coli* BL21 (DE3) with 15 ng of plasmid to generate the engineered cell without *xr*. The transformed cells were selected by growing on LB with 100 µg/ml ampicillin. The pETDuet-*luxCDEAB* plasmid and pRSFDuet-*gst*-*xr* plasmid were co-transformed into competent *E. coli* BL21 (DE3) with 15 ng of each plasmid to generate the engineered cell with *xr*. The transformed cells were selected by growing on LB with 100 µg/ml ampicillin and 34 for µg/ml kanamycin. A single colony was selected and grown in LB with appropriate antibiotic mentioned above as a starting liquid culture overnight at 37 °C. The corresponding variant was then sub-cultured into LB media with 1% inoculant (by volume) and grown at 37 °C until the OD_600_ reached 1.0. The culture was then induced with 10 mM lactose and incubated further at 25 °C for 24 h. Bacterial cells were harvested at designated times (0, 2, 4, 8, 12, 24 h) for measuring bioluminescent light. Relative Light Units (RLUs) were measured using a luminometer (AB-2270 Luminescencer-Octa, ATTO) and normalized with OD_600_ (RLU/OD_600_).

### Alkane whole-cell biocatalyst preparation and bioconversion assay

The pET28a-*fap* plasmid was transformed into competent *E. coli* BL21 (DE3) with 20 ng of plasmid to generate the engineered cell without *xr*. The transformed cells were selected by growth on LB with 34 µg/ml kanamycin. The pET28a-*fap* plasmid was transformed into competent *E. coli* BL21 (DE3), *glmS::pT7-gst-xr, Gm*^*r*^ with 20 ng of plasmid to generate the engineered cell with *xr*. The transformed cells were selected by growth on LB with 34 µg/ml kanamycin. A single colony was selected and grown in LB with the appropriate antibiotic mentioned above as a starting liquid culture overnight at 37 °C. The corresponding variant was then sub-cultured into TB media with 1% inoculant and grown at 37 °C until the OD_600_ reached 0.6. The culture was then induced with 75 mM lactose and incubated at 25°C for 20 hours in the dark. The bacterial cells were harvested by centrifugation and resuspended in 0.1 M Kpi buffer pH 7.0 to yield an OD_600_ of 60/mL. The cells were handled under red light. Bioconversion was done in 1 mL scale in a 20-mL clear glass capped vial containing 10 mM tetradecanoic acid as a substrate for the reaction. Reactions were illuminated under blue light (PPFD-B 20 µmolphotons/m^2^/s) and shaken at 100 rpm, 25°C. For alkane and fatty acid analysis, the reactions were extracted using ethyl acetate containing internal standard (200 µM tetradecane). The solvent layer was analyzed by Agilent 8890 GC-FID equipped with a HP5 column.

### Quenching of the samples for intracellular metabolites analysis

The method for quenching was performed following Winder *et al*, 2008 (*4*). Samples (1 ml) were rapidly quenched by adding equal volume of cold 60% methanol/water (-40 °C). The quenched samples were centrifuged for 10 min at 4°C and 800’*g*. The supernatant was removed, and the pellets were snap frozen in liquid nitrogen and stored at -80°C until metabolite extraction.

### Intracellular metabolite extraction

Intracellular metabolites were extracted using a cold-methanol method following Winder *et al*, 2008 (*4*). The pellets were resuspended in 2 mL of cold 100% methanol (-80 °C) (HPLC grade). The samples were frozen in liquid nitrogen and thawed on ice for 3 cycles for extracting intracellular metabolites. The suspensions were centrifuged at 18,800’*g* at -9°C for 30 min. The supernatant was transferred into a new 15 mL falcon tube, frozen in liquid nitrogen, and freeze-dried at -80°C, 1 mbar, for 16 hours. The dried samples were maintained on ice, resuspended in 500 µL of cold 50% ACN/water (HPLC grade), and centrifuged for 30 min at 18,800’*g* and at -9°C. The supernatant was collected and stored at -80 °C until analyzing with mass spectrometry.

### Mass spectrometry analysis

Samples (3 µL) were injected into an ion mobility drift tube-quadrupole time-of-flight mass spectrometer (LC-IM-QTOF, Agilent 1290 series LC 6560). An Agilent Poroshell 120 HILIC-Z 150 × 2.1 mm, 2.7 µm (particle size) column was used to achieve optimal separation of metabolites. The flow was 0.3 ml/min with a mobile phase of 10 mM ammonium acetate in water pH 9.0 containing 2.5 µM deactivator (Mobile phase A) and 10 mM ammonium acetate in 85% ACN pH 9.0 containing 2.5 µM deactivator (Mobile phase B). The gradient was changed from 4% of mobile phase A / 96% of mobile phase B to 35% of mobile phase A / 65% of mobile phase B in 24 min and the column was maintained at 35°C. Data acquisition was performed in IM-QTOF mode. The mass spectrometer was operated in negative ion mode. MS parameters were as follows: gas temperature, 250 °C; sheath gas temperature, 300°C; sheath gas flow, 12 l/min; fragmentor, 350 V; nozzle voltage, 1000 V; nebulizer, 45 psi; Vcap, 3000 V. The TOF mass was set as *m/z* 50-1700. For ion mobility parameters, high pure nitrogen (N_2_) was used for the drift gas. Other IM parameters were set as follows: IM-MS acquisition rate, 1 frame/s and 19 IM transients/frame; entrance and exit voltages of drift tube, 1600 and 250 V; trap filling and trap release times, 3200 and 250 μs; the drift tube pressure, 3.95 Torr. The drift time was limited to not more than 50 ms. The CCS values were calculated with the single electric field method. All data acquisitions were carried out using MassHunter Workstation Data Acquisition Software (Version B.08.00, Agilent Technologies, USA).

### Data processing and metabolite annotation

Data obtained from LC-IM-QTOF were preprocessed following Zhou et al, 2020 (*5*). Raw MS data files (.d) were first demultiplexed using the Agilent De-multiplexing tool (Version 1.0, Agilent Technologies). The demultiplexing data files (.DeMP.d) were recalibrated using IM–MS Reprocessor (Version B.08.00, Agilent Technologies). The reference masses used for mass calibration were m/z 112.985587 and 1033.988109. The CCS calibration was performed by IM–MS Browser software (Version 10.0, Agilent Technologies). The preprocessed data files were submitted for feature finding, alignment, and mass spectra extraction using Mass Profiler (Version 10.0, Agilent Technologies). Finally, the peak table and mass spectra (CEF format) files were exported for statistical analysis and metabolite annotation using Mass profiler professional (Version 15.1, Agilent Technologies) and ID browser identification (Version 10.0, Agilent Technologies), respectively. The detailed parameters of data processing tools were provided in Supplementary Table **5**. The metabolites were annotated using the parameters of accurate mass, isotope ratios, abundances and spacing, CCS, and retention time matching to standards. The *m/z* and CCS tolerance were set at 35 ppm and 5%, respectively. [M-H]^-^, [M+CH_3_COO]^-^, [M-2H]^2-^, [2M+CH_3_COO] ^-^, and [2M-H]^-^ adducts were considered for negative modes. The metabolite databases (METLIN, AllCCS (*5*), HMDB (*6*), MetCCS (*7*)) were used for known and unknown metabolite annotation.

### Statistical analysis

The data files in CEF format were imported to MPP for statistical analysis and visualization. Three separate projects (fatty alcohol, bioluminescence light, and alkane) were created in MPP. Under each project the data files were log_2_ transformed, normalized to total abundance of all samples, and baselined to the median of all samples. The features found in the data file were filtered based on frequency (Table **S5**) and on sample variability (Coefficient of variation ≥ 20 %). Unsupervised principal component analysis (PCA) was performed with mean centering and scaling to display the variance between the two groups (two strains of engineered cells: with and without XR). Statistical evaluation of the data was performed using univariate analyses(*8*). A cutoff value of *p* < 0.05 was considered statistically significant in unpaired t-test, using the Benjamini and Hochberg False Discovery Rate set to 5% for multiple testing corrections.

### Intra- and extra-cellular sugar analysis

Concentrations of lactose, glucose, galactose, sorbitol, and galactitol were measured overtime during the bioconversion to produce fatty alcohol (both intracellular and those excreted into the buffer solution). For intracellular sugar analysis, cell pellet was resuspended in water and lysed by ultrasonication. Proteins and cell debris were precipitated with 0.15% formic acid and centrifuged at 18,800ξ*g* at 4°C for 20 min. The resulting supernatant was filtered by a membrane with pore size 0.22 µm and kept at -20°C until further analysis. Sugars were identified and measured by LC-triple quadrupole mass spectrometry Agilent 1200 series LC 6470. The mass spectrometer was operated in a negative SIM mode to detect sugar and sugar alcohol based on the parameters of retention time and *m/z* by comparison to the values of standard compounds. MS parameters were as follows: gas temperature, 250°C; gas flow, 12 l/min; nebulizer, 45 psi; sheath gas flow, 12 l/min; capillary voltage, 3000 V; and VCharging, 1000. Data are shown as mean ± s.d., n = 3 replicate cultures; error bars show s.d; asterisk denotes significant differences by multiple t-test (*p* < 0.05). The Agilent Hi-plex H 300 × 7.7 mm column operated at 35°C was used to achieve optimal separation. The flow rate was 0.3 mL/min using an isocratic mobile phase of 0.1% formic acid in water.

### Gene expression analysis by quantitative real-time PCR (qRT-PCR)

To identify the up-regulated genes in the engineered XR-harboring cells in accordance with untargeted metabolomics analysis, the fatty alcohol biocatalyst during the bioconversion process was subjected to analysis of gene expression. Total RNA was extracted from the cell pellet by hot acid-phenol/chloroform method and converted into complementary DNA (cDNA) by reverse transcription. The qRT-PCR was performed using KAPA SYBR FAST qPCR Master Mix (2X) Kit (KAPA Biosystems) with StepOnePlus Real-Time PCR system (Applied Biosystems). The gene expression level was calculated by *ΔΔ*Ct method normalized with the *16s rRNA* gene and compared relative to those from the reference sample. Primers for PCR were listed in Table **S7**. Multiple unpaired t-test was used to compare a mean of each target gene from different strains by GraphPad Prism version 9.3.1 (GraphPad Software, San Diego, California USA). The cutoff value of *P* < 0.05 was adjusted for multiple comparisons using Bonferroni-Dunn method and considered as statistically significant different.

## Author Contributions can be described as follow

conceptualization: JJ, CS, PC; Methodology: JJ, CS, JP, WC, CK, NA, NK, KP, SA, MF, PC; Investigation: JJ, CS, JP, WC, CK, NA, NK, KP, SA; Resources: TP, FH, MF, PC; Supervision: PC; Writing – Original Draft: JJ, CS, PC; Writing – Review & Editing: JJ, CS, PC.

## Supplementary Materials

**Fig. S1.**
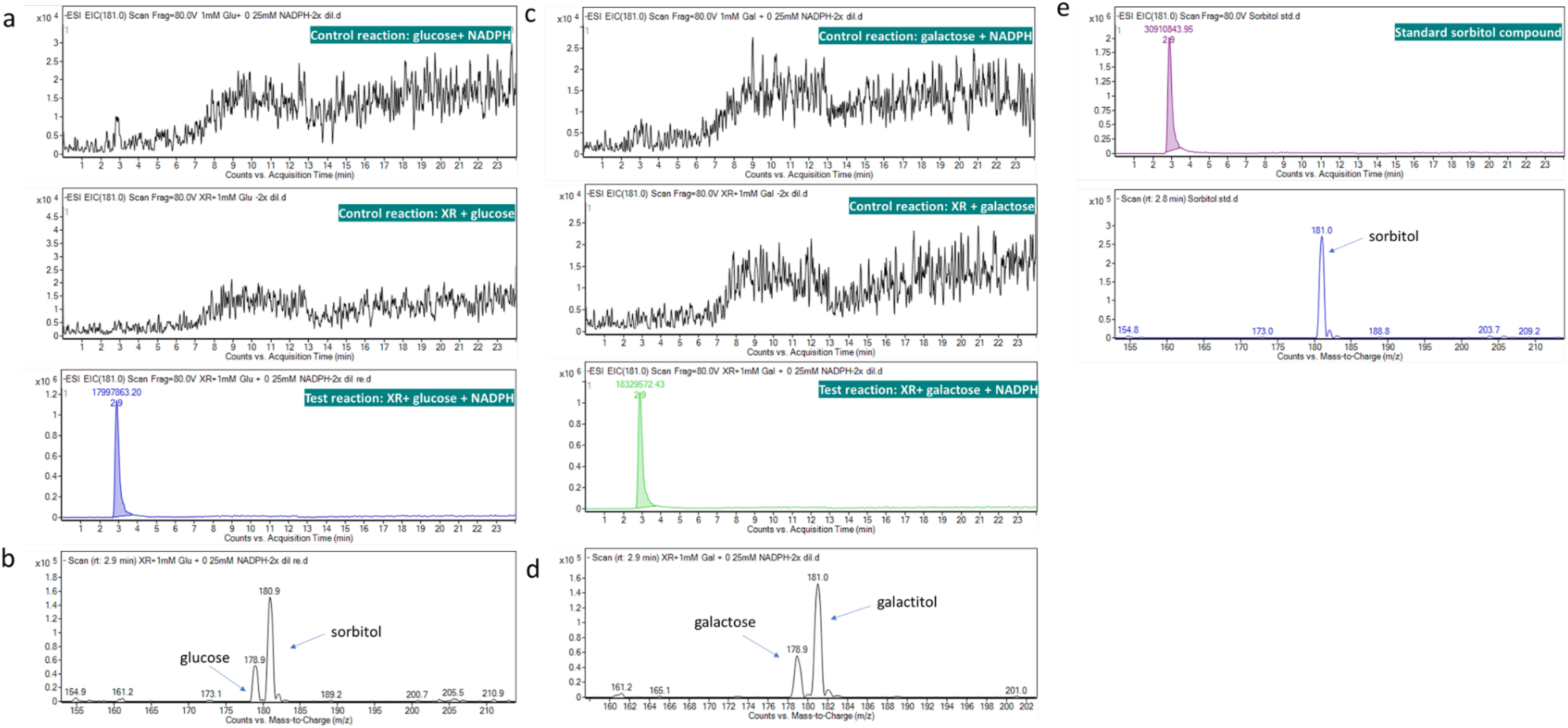
Sugar reduction by XR using D-glucose and D-galactose as substrates. **(a**-**b)** The extracted ion chromatograms (EIC) and exact mass spectra at *m/z* 181.0 ([M-H]^-^) identified sorbitol as a product from reduction of D-glucose by XR. The reactions were carried out under ambient temperature for 10 min in 100 mM potassium phosphate buffer, pH 7.5 containing 5 µM XR, 0.25 mM NADPH, and 1 mM glucose. The control reaction was performed without addition of XR and NADPH. The sorbitol product was analyzed by an LC-MS triple quadrupole mass spectrometer in negative mode. **(c**-**d)** The extracted ion chromatogram (EIC) and exact mass spectra at *m/z* 181.0 ([M-H]^-^) identified galactitol as a product from the reduction of D-galactose by XR. The reactions were carried out under ambient temperature for 10 min in 100 mM potassium phosphate buffer, pH 7.5 containing 5 µM XR, 0.25 mM NADPH, and 1 mM galactose. The control reaction was performed without addition of XR or NADPH. The galactitol product was analyzed by LC-MS with a triple quadrupole mass spectrometer in negative mode. **(e)**The extracted ion chromatogram (EIC) and exact mass spectra at *m/z* 181.0 ([M-H]^-^) of a standard sorbitol analyzed by an LC-MS triple quadrupole mass spectrometer in negative mode. The standard compound and samples were diluted in acetonitrile (ratio 1:1) before being analyzed on an Agilent 1200 series LC 6470 triple quadrupole mass spectrometer. The Agilent Poroshell 120 HILIC-Z 150 × 2.1 mm, 2.7 µm (particle size) column was used to achieve optimal separation. The mobile phase was 10 mM ammonium acetate in water pH 9.0 containing 2.5 µM deactivator (mobile phase A) and 10 mM ammonium acetate in 85% ACN pH 9.0 containing 2.5 µM deactivator (mobile phase B) with a flow rate of 0.3 ml/min. The gradient was changed from 4% mobile phase A / 96% mobile phase B to 35% mobile phase A / 65% mobile phase B in 24 min, and the column was maintained at 35°C. The mass spectrometer was operated in scan mode to detect sugar alcohol products based on their retention times and *m/z* values compared to standard compounds. MS parameters were as follows: gas temperature, 250 °C; gas flow, 12 l/min; nebulizer, 45 psi; sheath gas flow, 12 l/min; capillary voltage, 3000 V; and VCharging, 1000.

**Fig. S2.**
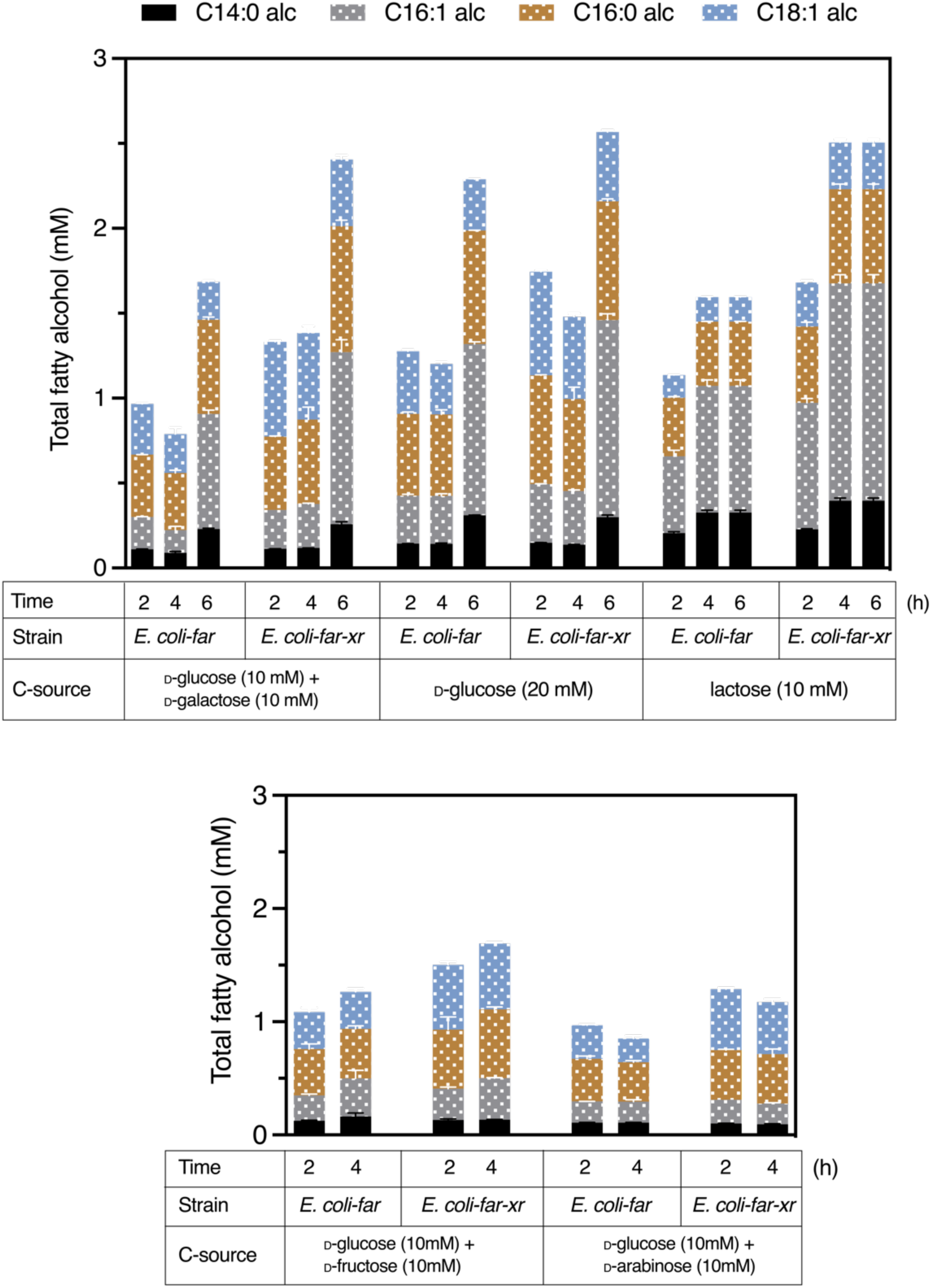
Amount of fatty alcohol (mM) produced by *E. coli*-*far* and *E. coli*-*far-xr* at various time points of bioconversion using various types of sugars. *E. coli*-*far* and *E. coli*-*far-xr* were induced with 1 mM lactose at 220rpm, 25 °C for 6 h and then harvested and employed as biocatalysts for fatty alcohol production. The bioconversion reactions were carried out in potassium phosphate buffer containing cells with OD_600_ of 30/mL and 10 mM each of D-glucose/D-galactose, 20 mM D-glucose, 10 mM lactose, 10 mM each of D-glucose/D-fructose, and 10 mM each of D-glucose/D-arabinose. The reactions were incubated at 220 rpm, 25°C for 2, 4, and 6 h with 10 mM each of D-glucose/D-galactose, 20 mM D-glucose, and 10 mM lactose, and for 2 to 4 h for 10 mM each of D-glucose/D-fructose and 10 mM each of D-glucose/D-arabinose. Fatty alcohol products were extracted with 2 mL of ethyl acetate containing internal standard (200 µM of tetradecane). Data are shown as mean ± s.d., n = 2 replicate cultures. The addition of XR clearly improves the yield of fatty alcohol production by the whole-cell biocatalysts with all tested sugars, particularly with mixed D-glucose/D-galactose and lactose utilization. This might be due to XR conversion of glucose into its corresponding alcohol (sorbitol), alleviating the glucose repression effects and allowing the cell to co-utilize other carbon sources for creating the necessary precursors simultaneously for fatty alcohol synthesis. As glucose is generally preferred by microbes for use in its metabolism, the addition of XR did not cause as much of a profound effect as the presence of multiple carbohydrate substrates. The addition of XR thus offers an alternative method for improving the utilization of multiple carbohydrate substrates.

**Fig. S3.**
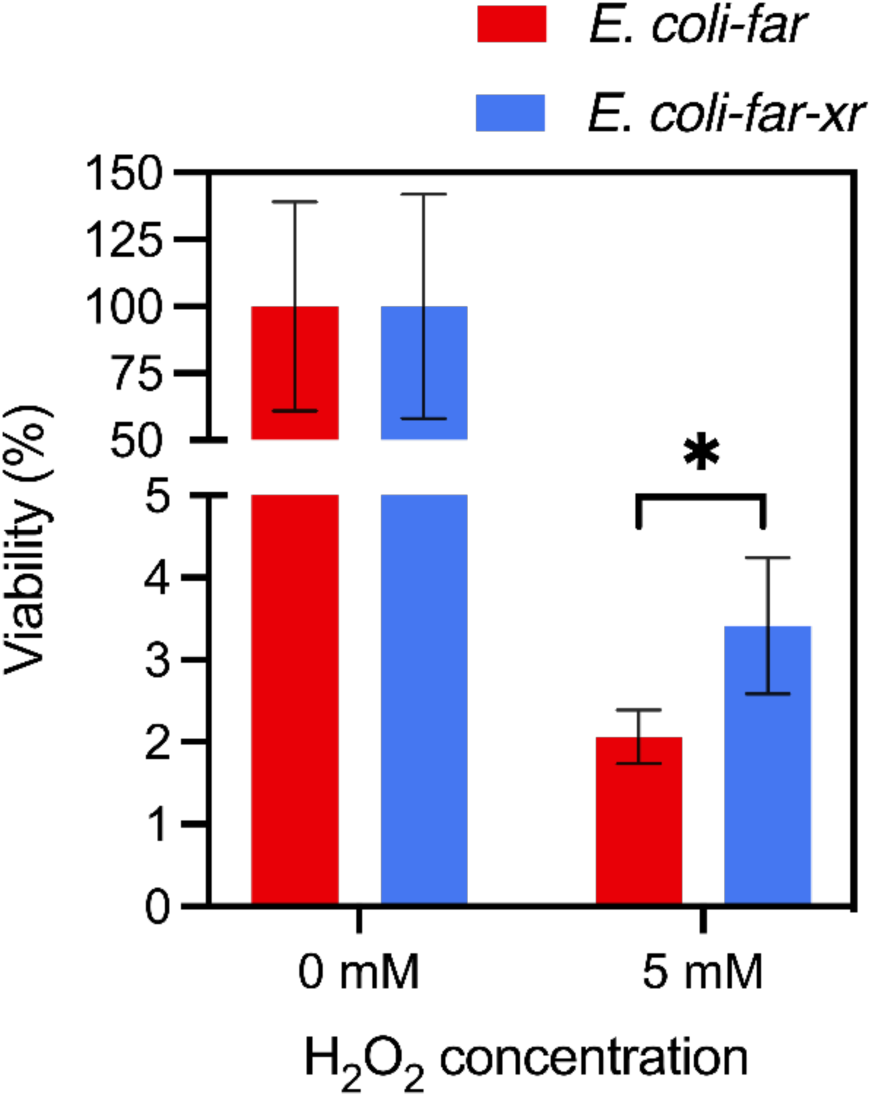
H_2_O_2_ susceptibility test on *E. coli-far* and *E. coli-far-xr*. *E coli* strain FAR and FAR-XR protein overexpression was induced with 1 mM lactose for 6 h and used for an oxidative stress assay by adding H_2_O_2_. A 0.5-mL reaction contained the biocatalyst with final OD_600_ of 0.4/mL and 5 mM H_2_O_2_ in 0.1 M Kpi pH 7.5. After incubation for 90 min at room temperature, the viable cells were measured by colony forming unit (CFU) counting on an LB agar plate. The results suggested that the cells containing XR had higher CFUs than the cells without XR. Asterisk indicates significant differences by *t* test (*P* < 0.05).

**Fig. S4.**
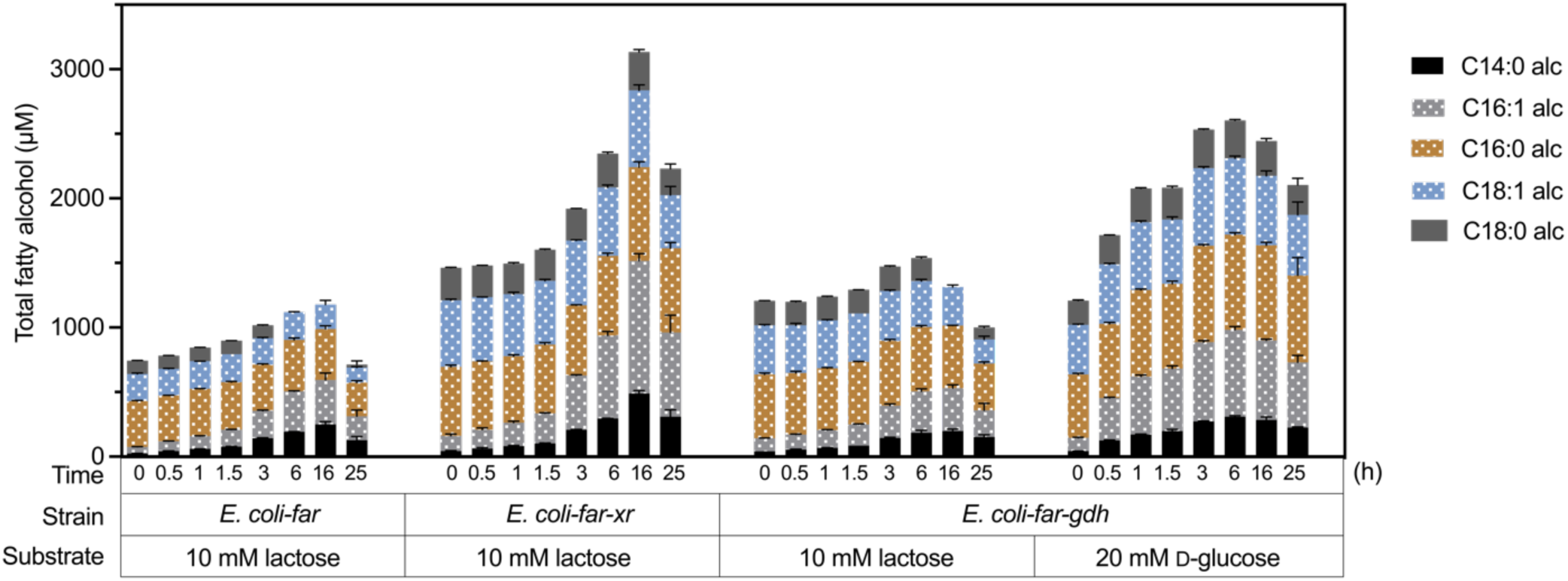
Comparison of fatty alcohol production from different cofactor regenerating systems in the engineered cell *E. coli*-*far*. *E. coli*-*far-gdh* cells were tested with both using 10 mM lactose and 20 mM D-glucose as substrates for generation of sugar phosphates for fatty alcohol production. The bioconversion and fatty alcohol detection were described in Supplementary Methods. Data are shown as mean ± s.d., n = 3 replicate cultures.

**Fig. S5.**
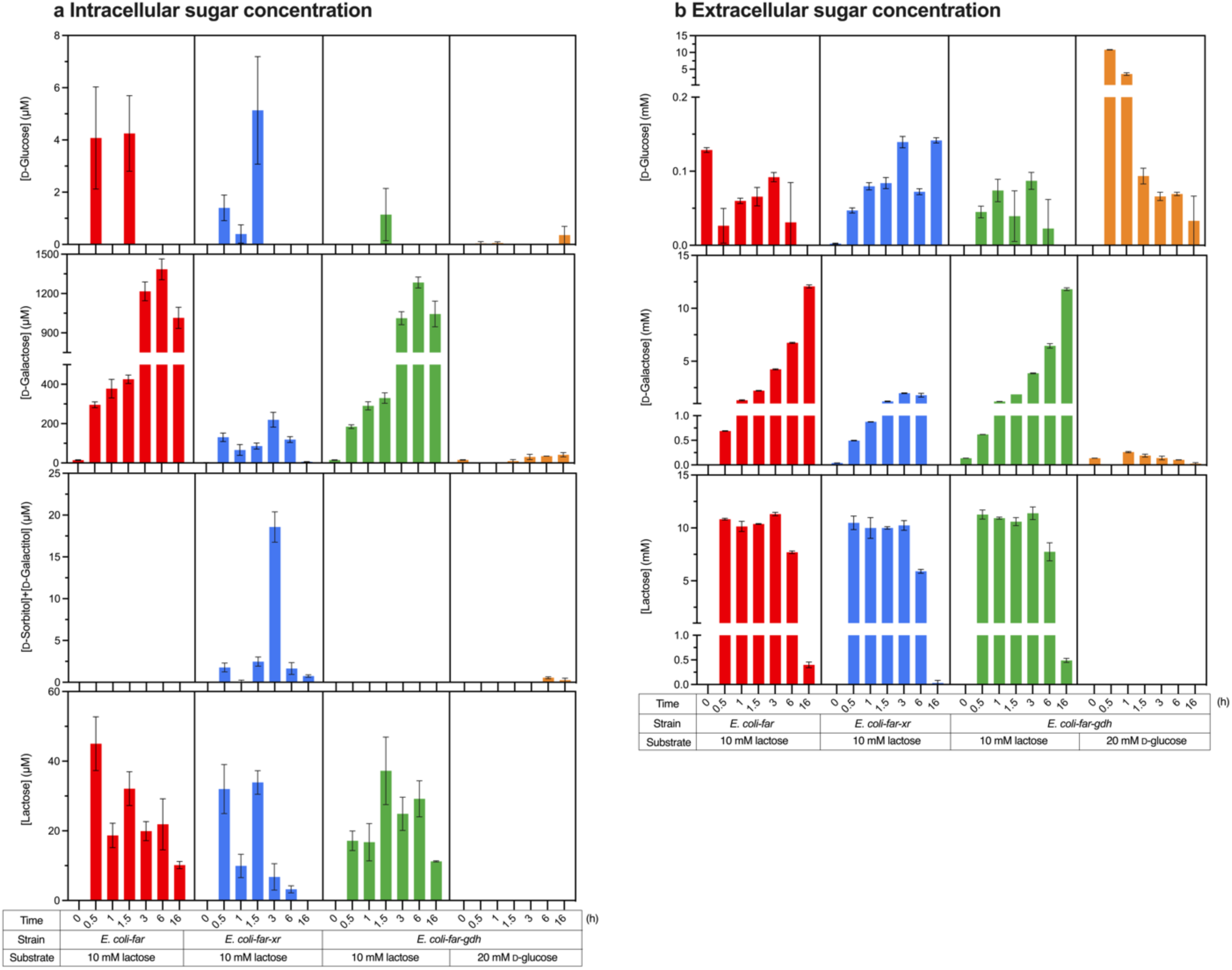
Measurement of sugars from the fatty alcohol producing cells. Sugars were detected during the bioconversion in *E. coli-far* (red), *E. coli-far-xr* (blue), *E. coli*-*far-gdh* when 10 mM lactose was used as a substrate (green) and when 20 mM D-glucose was used as a substrate (yellow) in *E. coli*-*far-gdh*. The bioconversion and sugar detection protocols were described in the Supplementary Methods. These graphs show the sugars from both inside the cell (**a**) and those secreted into media (**b**) over time. Sorbitol/galactitol was not found in extracellular sugars in all samples. Noted that samples from Time 0 h were those before sugars were added into the reactions. Analytical methods were described in the Methods section. Data are shown as mean ± s.d., n = 3 replicate cultures.

**Fig. S6.**
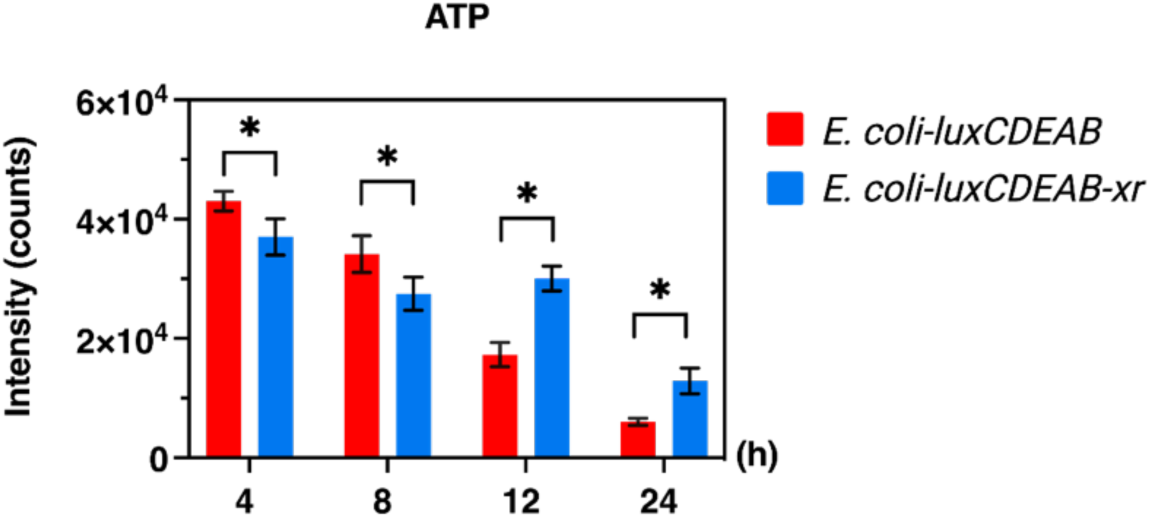
Time-course analysis of ATP in *E. coli*-*luxCDEAB* and *E. coli*-*luxCDEAB-xr* after adding 10 mM lactose for 4, 8, 12, and 24 h. The samples were diluted in acetonitrile (ratio 1:1) before being analyzed by the LC 6470 triple quadrupole mass spectrometer (Agilent 1200 series). An Agilent Poroshell 120 HILIC-Z 150 × 2.1 mm, 2.7 µm (particle size) column was used to achieve optimal separation. The flow was 0.3 ml/min with a mobile phase of 10 mM ammonium acetate in water pH 9.0 containing 2.5 µM deactivator (mobile phase A) and 10 mM ammonium acetate in 85% ACN pH 9.0 containing 2.5 µM deactivator (mobile phase B). The gradient was changed from 4% of mobile phase A / 96% of mobile phase B to 35% of mobile phase A / 65% of mobile phase B over 24 min., and the column was maintained at 35°C. The mass spectrometer was operated in a SIM mode to detect ATP based on the parameters of retention time and *m/z* by comparison to values of the standard compound. MS parameters were as follows: gas temperature, 250°C; gas flow, 12 l/min; nebulizer, 45 psi; sheath gas flow, 12 l/min; capillary voltage, 3000 V; and VCharging, 1000. Data are shown as mean ± s.d., n = 5 replicate cultures; error bars show s.d; asterisk denotes significant differences by multiple *t* test (*P* < 0.05).

**Fig. S7.**
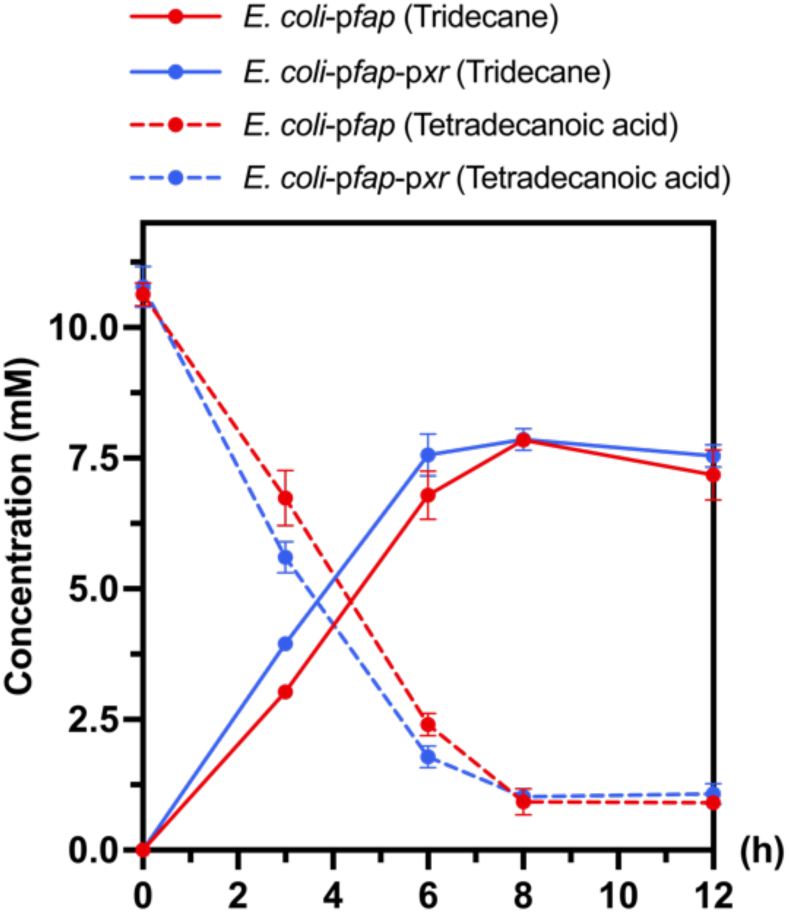
Tridecane production (mM) from tetradecanoic acid by *E. coli* harboring FAP with/without XR overexpressed in the plasmid system. Plasmids of pCDFDuet-*fap* and pRSFDuet-*gst*-*xr* (pRSFDuet in case of biocatalyst without XR) were co-transformed into *E. coli* BL21 (DE3) to generate *E. coli*-p*fap* and *E. coli*-p*fap*-p*xr*. A single colony of the resulting transformants was inoculated in LB broth in the presence of streptomycin 25 µg/mL and kanamycin 34 µg/mL at 220 rpm, 37°C for 17 h. The starter cultures were then sub-cultured with 1% inoculant into TB media containing a half-dose of the antibiotics mentioned above and grown at 37 °C until the OD_600_ reached 0.6-0.7. The cultures were then induced with 10 mM lactose and incubated further with shaking at 220 rpm, 25 °C for 12 h, then harvested and employed as biocatalysts for alkane production. The bioconversion reactions were carried out in potassium phosphate buffer (0.1 M, pH 7.0) containing a cell biocatalyst at an OD_600_ of 60/mL and 10 mM tetradecanoic acid. The reaction was set in a capped 20mL clear glass vial and incubated under blue light (PPFD-B 20 µmolphotons/m^2^/s) with shaking at 100 rpm, 25°C. Fatty acid and alkane were extracted using 4 mL of ethyl acetate containing the internal standard (200 µM of tetradecane) and analyzed by GC-FID equipped with HP-5 column. Data are shown as mean ± s.d., n = 3 replicate cultures. 3 hours after the reaction was initiated, the productivity of *E. coli*-p*fap*-p*xr* (blue) was 1.3 mmol/L/h, which is higher than 1 mmol/L/h of *E. coli*-p*fap* (red) by 1.3-fold. To improve the use of the XR/lactose system as an enhancer system for producing the active FAP (FAP:FAD) without adding extra antibiotic and creating a more stable biocatalyst, we further created a strain of *E. coli* BL21 (DE3) with an XR integrated genome (see construction details in Supplementary Methods). The use of the XR genome integrated cell version for production of tridecane was discussed in the Main text.

**Fig. S8.**
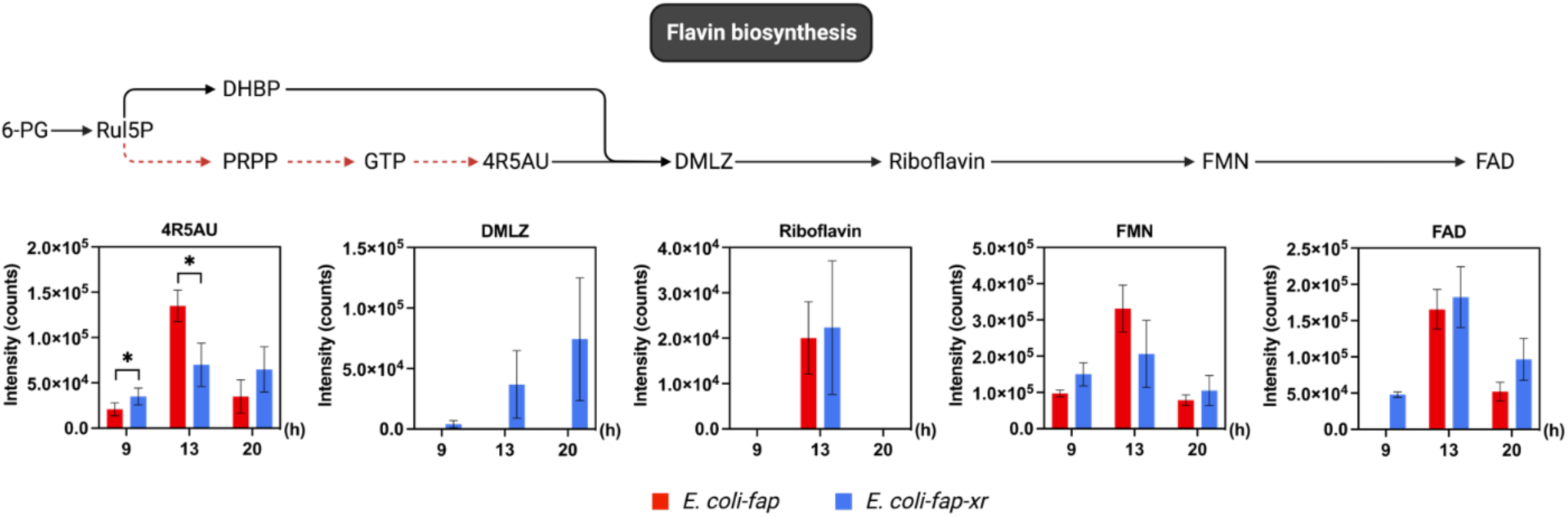
Level of intermediates in flavin biosynthesis during protein overexpression of *E. coli*-*fap* and *E. coli*-*fap-xr*. After 75 mM of lactose was added into the TB culture of *E. coli*-*fap* and *E. coli*-*fap-xr*, the cells (at equivalent cell amounts based on OD_600_ in 1 mL) were taken from the culture at 9, 13, and 20 hours. The metabolites were extracted from the samples and analyzed by LC-IM-QTOF as described in the Methods. The levels of metabolites were obtained from the peak area by Find by the Formular (FBF) function with mass tolerance ±30 ppm in the MassHunter Qualitative software, with the exception of DMLZ, for which levels were obtained from the abundance value as identified by Mass Profiler. Data are shown as mean ± s.d., n = 6 technical replicates (2 biological replicates), and the asterisk denotes a significant difference by multiple *t* test (*P* < 0.05, Bonferroni-Dunn method).

**Fig. S9.**
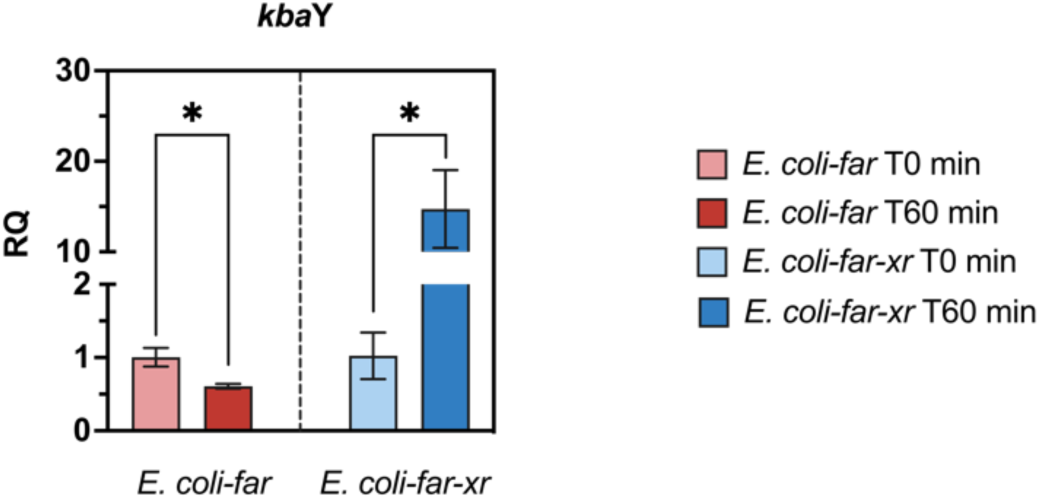
Expression levels of sorbitol/galactitol degradation genes (*kbaY*) in *E. coli*-*far* and *E. coli*-*far-xr*. Analysis of real-time quantitative PCR of samples from T0 min and T60 min during the fatty alcohol bioconversion process. Y-axis represents relative quantification (RQ) of each biocatalyst at T60 min relative to its level at T0 min. Data are shown as mean ± s.d., n = 3 replicate cultures. The asterisk denotes a significant difference by *t* test (*P* < 0.05).

**Fig. S10.**
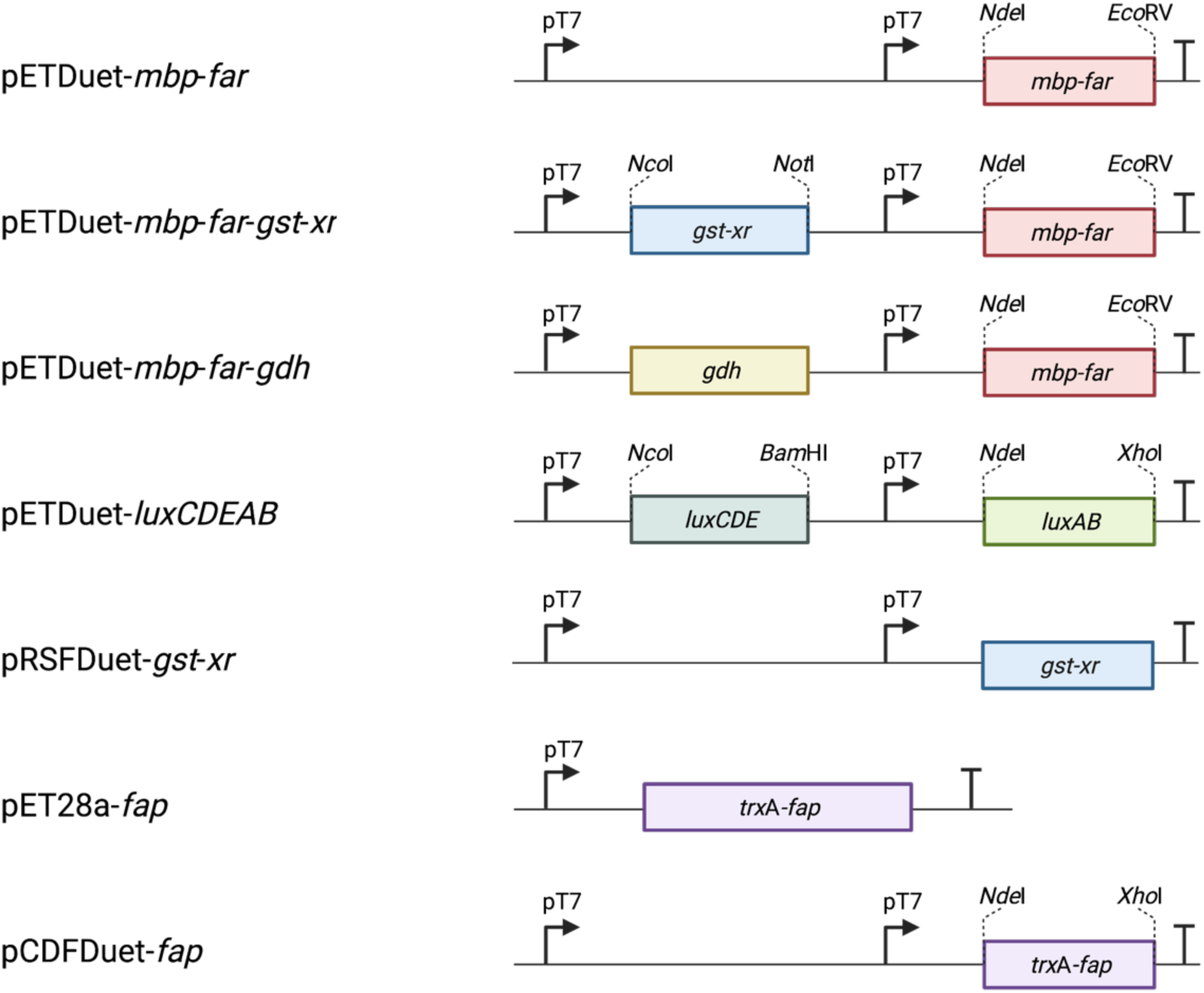
Expression plasmid design used in this study. Details of construction are described in Supplementary Methods; pT7 = T7 promoter.

**Data for Fig. S2.**
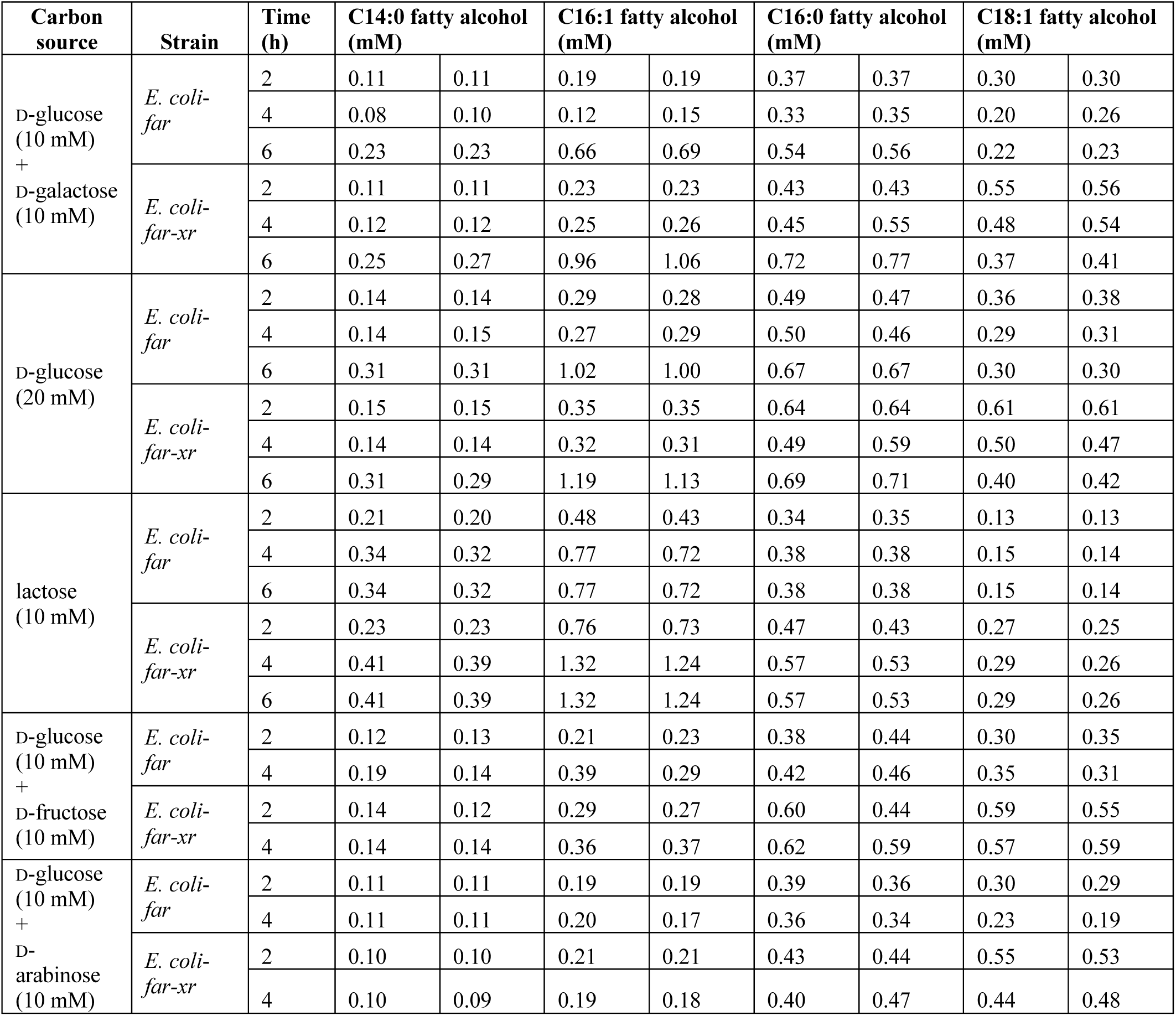
Amount of fatty alcohol (mM) produced by *E. coli-far* and *E. coli*-*far-xr* at various time points of bioconversion using various types of sugar.

**Data for Fig. S3.**
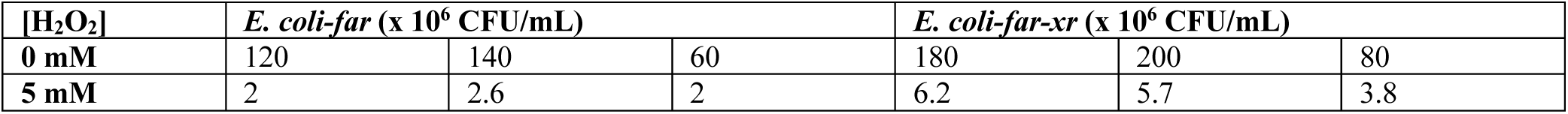
Colony forming unit (CFU) counting on an LB agar plate.

**Data for Fig. S6.**
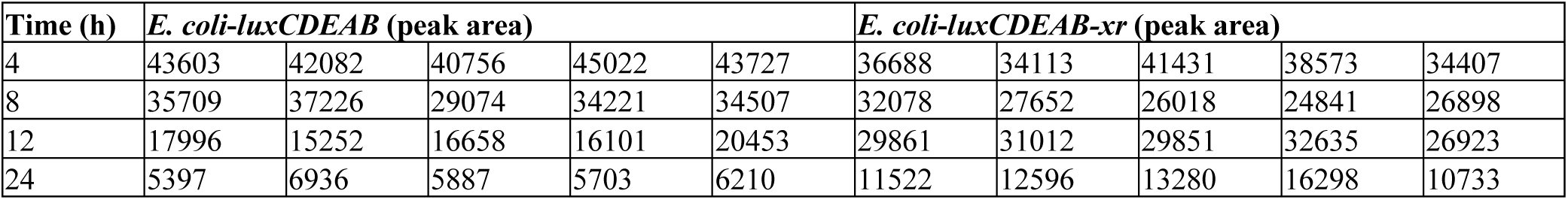
Time-course analysis of ATP in *E. coli*-*luxCDEAB* and *E. coli*-*luxCDEAB-xr* after adding 10 mM lactose for 4, 8, 12, and 24 h.

**Data for Fig. S7.**
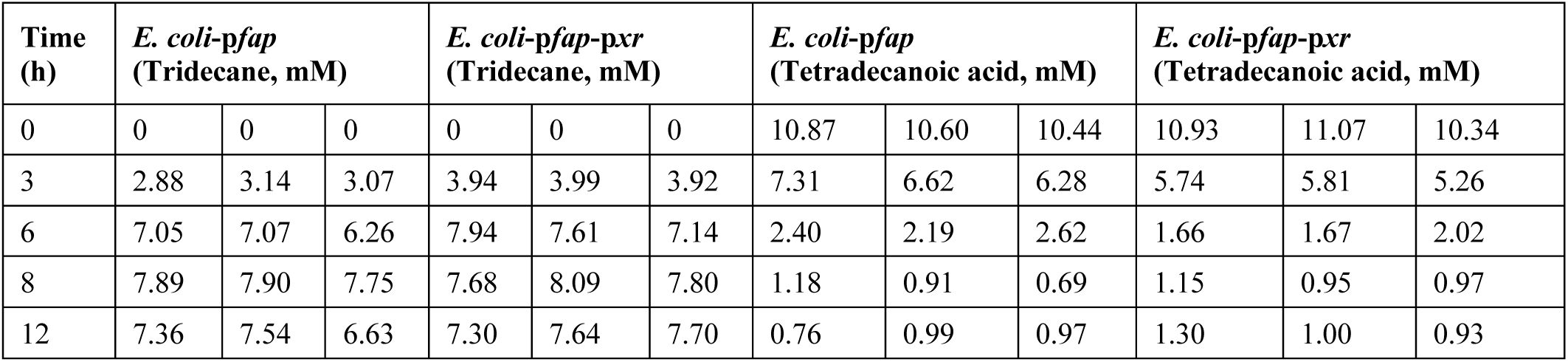
Tridecane production (mM) from tetradecanoic acid by *E. coli* harboring FAP with/without XR overexpressed in the plasmid system.

**Data for Fig. S8.**
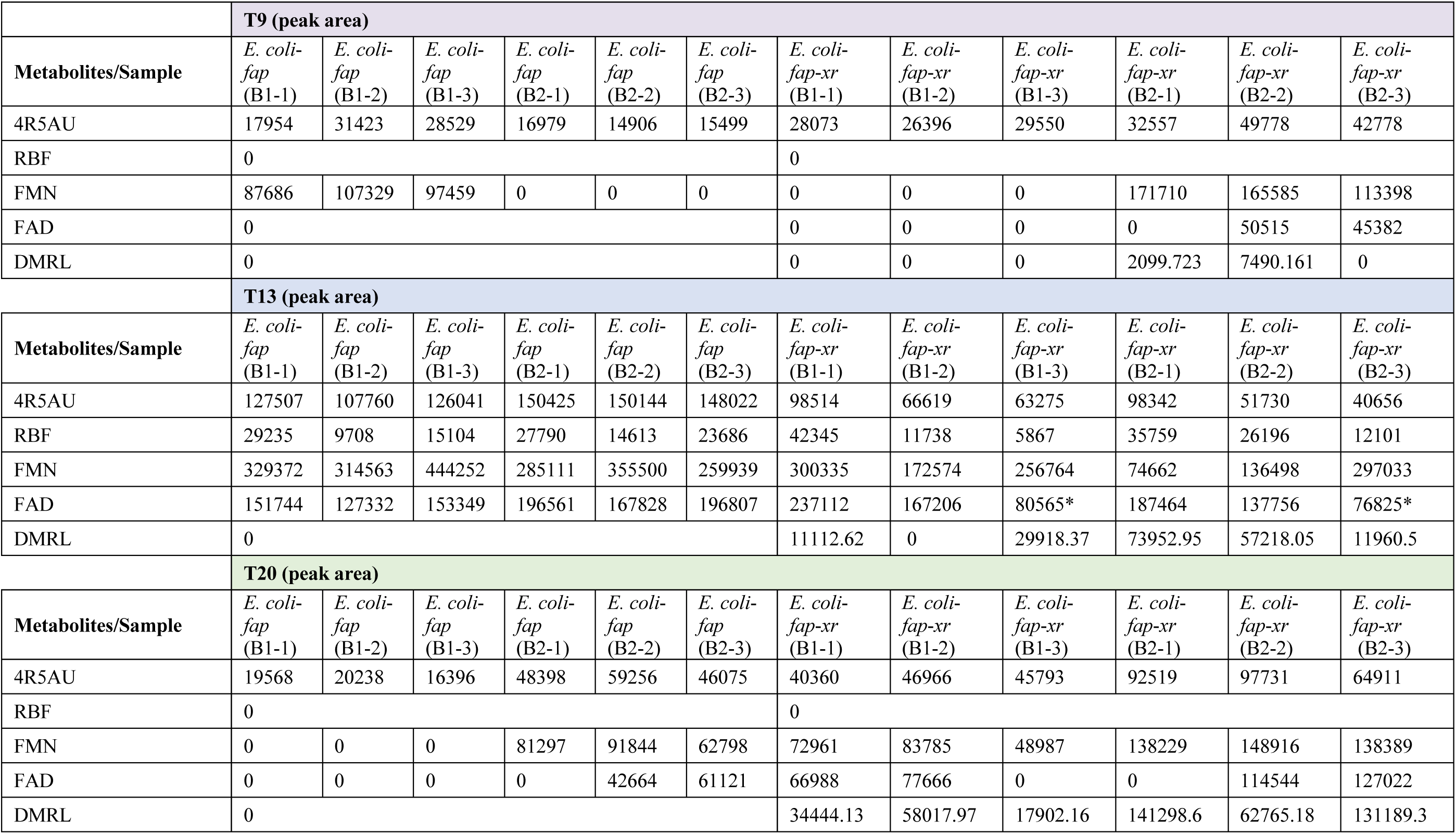
Levels of intermediates in the flavin biosynthesis during protein overexpression of *E. coli*-*fap* and *E. coli*-*fap-xr*. *Data points were excluded from the graph based on technical outliers.

Table S1. The annotated metabolites of fatty alcohol biosynthesis

Table S2. The annotated metabolites of bioluminescense light generation

Table S3. The annotated metabolites of alkane biosynthesis

Table S4. Primers used for plasmid construction

Table S5. Parameters used for data processing

Table S6. Abbreviations

Table S7. Primers used for qRT-PCR

## Footnotes

1 Note that while specialty compounds such as NAD(P)H, NAD(P)^+^, ATP and acetyl-CoA have roles as substrates in most reactions, because they bind to enzymes in particular (redox) forms and dissociate from enzymes in other forms, they are often referred to in the literature as cofactors or coenzymes because of their re-current roles in various reactions in metabolic pathways. Although FAD and FMN mostly bind constitutively to enzymes, substantiating their designation as cofactors, they can also serve as substrates by having different redox forms bind to and dissociate out from particular enzymes such as in two-component flavin-dependent monooxygenase (*44, 45*). To simplify the terminology used, this work refers to compounds (NAD(P)H, NAD(P)^+^, FMN, FAD, acetyl-CoA) recurrently found in metabolic pathways as cofactors.

